# Regulated Cleavage and Relocation of FERONIA Control Immunity in Arabidopsis Roots

**DOI:** 10.1101/2023.10.16.562456

**Authors:** Jia Chen, Xiaonan Qiang, Hongbin Liu, Fan Xu, Long Wang, Lingli Jiang, Chiyu Li, Bingqian Wang, Sheng Luan, Dousheng Wu, Feng Zhou, Feng Yu

## Abstract

Plant roots exhibit localized immunity (LI) specifically in the root cap and transition/elongation zone (TZ/EZ). Such immunity relies on receptor-like kinases localized at the plasma membrane (PM), which are essential for plant response to bacteria in the rhizosphere. Here, we identified a new mechanism of action for receptor kinase FERONIA (FER) in LI. In the absence of bacterial stimuli, FER is localized to the PM, maintaining plant growth. When confronted with bacterial colonization, RALF23 peptide hyper-accumulates in TZ/EZ cells and activates a metalloproteinase that cleaves FER and transfer the cytosolic kinase domain into the nucleus. The nucleus-localized kinase domain of FER contributes to LI by reducing bacterial abundance at TZ/EZ. Thus, this work uncovered a unique mechanism by which a single receptor kinase acts in two molecular forms and two distinct subcellular locations to balance growth and immunity.

**One Sentence Summary:** Bacteria induces the cleavage of FER via At2-MMP to trigger a localized immunity around TZ and EZ of Arabidopsis roots.

## Introduction

Some members of the receptor-like kinases (RLKs) superfamily, serve as plasma membrane (PM) pattern recognition receptors (PRRs) to perceive microbe-associated molecular patterns (MAMPs) or host-derived damage-associated molecular patterns (DAMPs), leading to immune responses ^1,2^. Some RLK members orchestrate the trade-off between growth and immunity in response to endogenous plant signaling peptides^1,2^. Plant roots, similar to the human gastrointestinal tract, interact with countless microbes and must distinguish between those that pose a threat and those that offer potential benefits ^3–5^. Precise regulation of the on-and-off of immune responses is critical to ensure plants’ growth and prevent excessive defense responses in the absence of infection ^6,7^. Nevertheless, plants selectively protect vulnerable areas such as the transition and elongation zone (TZ and EZ), where commensal and pathogenic bacteria often colonize, to limit defense response in specific regions of roots ^8–10^. This phenomenon is referred to as localized immunity (LI). Because division and elongation of TZ and EZ, respectively, are also crucial for root growth, activation of LI would inhibit growth. Consequently, LI is important in balancing defense and growth ^9–13^, but the mechanisms how plants sense bacterial colonization to achieve LI remain unclear ^8^. Small peptides from the rapid alkalinization factors (RALFs) family alkalinize extracellular pH and inhibit root growth ^14,15^. Among them, RALF23 is unique in which it restrains immunity in leaves ^16,17^ through binding to its receptor FERONIA (FER) which belongs to the Catharanthus roseus receptor-like kinase (CrRLK1L) family and localizes to the plasma membrane (PM) ^18^. Intriguingly, genetic analysis indicates that FER has multiple and even opposite roles to RALF23 ^19–22^. FER is one of the best-studied RLKs in plants, but so far all of its functions have been attributed to its activity as a plasma membrane-localized receptor. In this study, we have identified a second form of FER produced through cleavage of its cytosolic domain that is then translocated into the nucleus. This cleavage and translocation occur in response to bacterial colonization that induces accumulation of RALF23 and subsequently activates a proteinase. The nucleus-localized FER will then activate immunity in cells around TZ and EZ of roots and thereby suppresses bacterial colonization.

## Results

### FER regulates localized immunity (LI) in roots

To gain insight into the function of FER in root immunity, we monitored the wild type and *fer-4* mutant ^23^ using the fluorescent MAMP marker lines, expressing a triple mVenus fused to a nuclear localization signal (NLS-3×mVenus) driven by the *PEROXIDASE 5* (*PER5*) or *FLG22-INDUCED RECEPTOR-LIKE KINASE 1* (*FRK1*) promoters ^9,24,25^. We first tested whether FER affects LI in roots in response to bacteria. We used the model commensal rhizobacterium *Pseudomonas protegens* strain CHA0 (CHA0) ^26,27^ and the bacterial pathogen *Pseudomonas syringae* pv. *tomato* DC3000 (DC3000) ^28^. CHA0 and DC3000 induced LI in WT (Fig. 1a-b). In comparison, LI response in the *fer-4* background was significantly reduced in response to CHA0 and DC3000 treatment (Fig. 1c-d). To test whether LI affects bacterial colonization of roots, we assessed root colonization of CHA0 and observed more growth in the TZ and EZ of *fer-4* than that in WT (Fig. 1e-f). Flagellin 22 (flg22), a well-described pathogen-associated molecular pattern (PAMP), is perceived by flagellin-sensing 2 (FLS2) to activate a series of downstream immune responses ^29,30^. Previous studies have shown that flg22 triggers a rapid and strong response in Arabidopsis roots mainly in TZ and EZ ^8–10^. Furthermore, a prolonged treatment with flg22 also induces an immune response in the meristematic zone (MZ) ^8,9^. Using flg22 as an immune elicitor, we found that flg22-triggered immunity was significantly reduced in the TZ and EZ of roots in *fer-4* mutant compared to the WT (Extended Data Fig.1a-d).

**Fig. 1.**
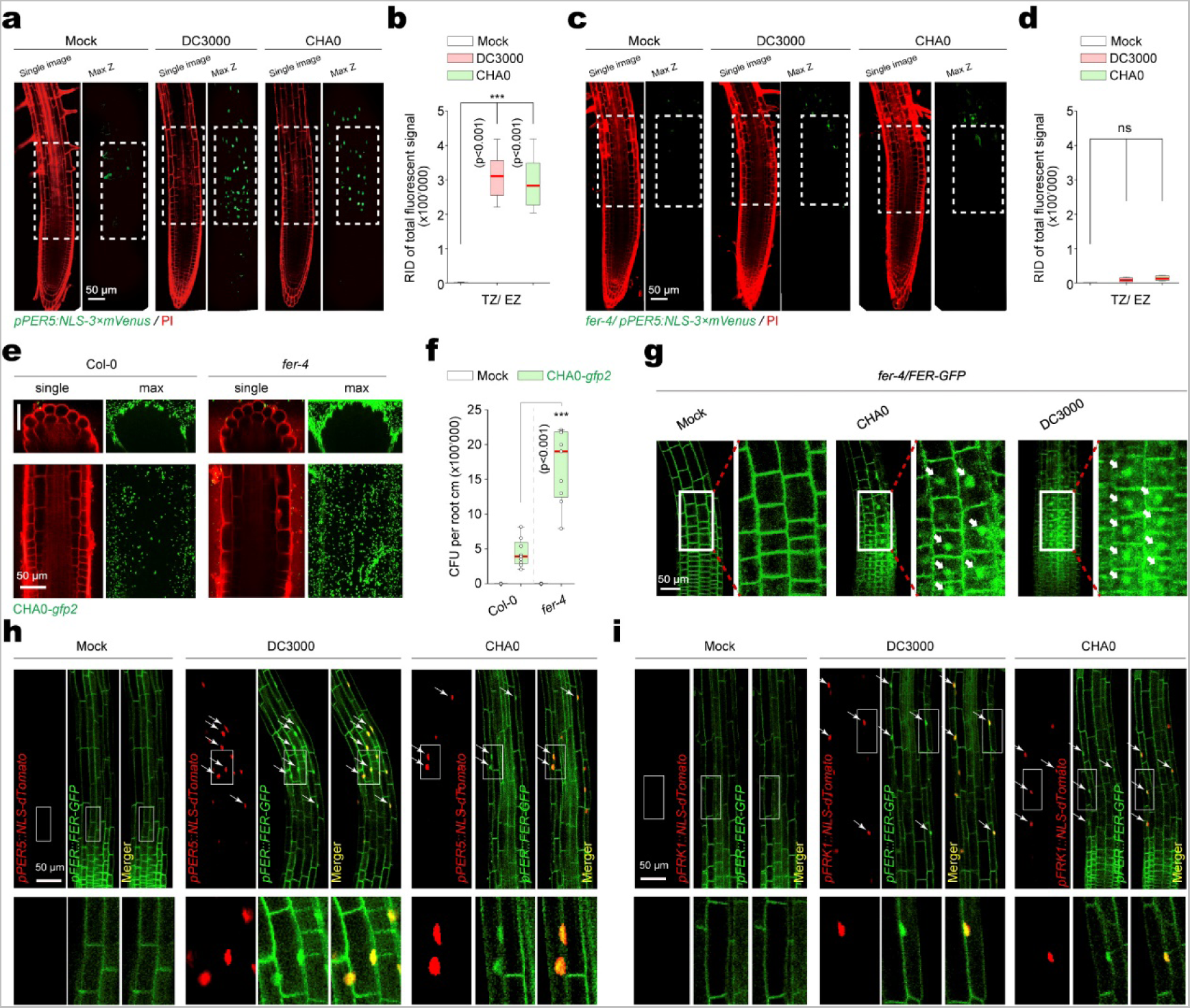
FER participates in localized immune responses around the TZ and EZ of roots. (**a**) mVenus fluorescence patterns in *pPER5::NLS-3×mVenus* line in the absence or presence (CHA0 at OD_600_ = 0.02 or DC3000 at OD_600_ = 0.1) of bacterial colonization for 18 h in the Col-0 or *fer-4* background. mVenus signals (green) co-visualized with propidium iodide (PI, red). Single confocal microscope image and maximal projections of Z-stacks are presented. (**b**) Quantitative analysis of mVenus signal intensities of the *PER5* marker in the absence or presence (CHA0 at OD_600_ = 0.02 or DC3000 at OD_600_ = 0.1). RID, raw intensity density. RID of total fluorescent signals in a single image is the sum of the RID of each nuclear signal in the imaged aera. Boxplot centers show median (n = 12 roots). Asterisks (***p < 0.001) indicate statistically significant differences between means by ANOVA and Tukey’s test analysis. (**c**) mVenus fluorescence patterns in *fer-4/ pPER5::NLS-3×mVenus* line in the absence or presence (CHA0 at OD_600_ = 0.02 or DC3000 at OD_600_ = 0.1) of bacterial colonization for 18 h in the Col-0 or *fer-4* background. (**d**) Quantitative analysis of mVenus signal intensities of the *PER5* marker in the absence or presence (CHA0 at OD_600_ = 0.02 or DC3000 at OD_600_ = 0.1). Boxplot centers show median (n = 12 roots). ns, not significant. (**e**) Representative images showing CHA0-*gfp2* colonization on Col-0, and *fer-4* roots in the TZ and EZ at 3 days post-inoculation (dpi). Photographs are maximum projections of confocal Z stacks. (**f**) Colony forming units (CFU) counting of CHA0-*gfp2* colonization on Col-0, and *fer-4* roots in the TZ and EZ at 3 dpi. Three roots were collected for each sample at the indicated colonization time point. Values are means ± SD (3 biological replicates). Asterisks (\*\*\**p* < 0.001) indicate significant differences based on ANOVA followed by Tukey’s test. (**g**) GFP fluorescence localization in D-roots of *fer-4/FER-GFP* in response to CHA0 at OD_600_ = 0.02 or DC3000 at OD_600_ = 0.1 treatments for 18 h in D-roots. (**h–i**) FERN co-localizes with PER5 (**h**) or FRK1 (**i**) in the TZ and EZ cells after treatment with CHA0 at OD_600_ = 0.02 or DC3000 at OD_600_ = 0.1 for 18 h. Arrows indicate the nucleus.

To confirm the role of FER in rhizobacteria-induced LI, we treated *fer-4/FER-GFP* ^23^ with CHA0 and DC3000. Intriguingly, GFP fluorescence was detected in the nucleus after CHA0 and DC3000 treatment in the TZ and EZ of roots (Fig. 1g). Moreover, the fluorescence signal associated with nuclear localization predominantly exhibited concentration within the epidermis (Extended Data Fig.1e). To examine if nuclear FER-GFP signal is associated with LI in response to bacteria, we co-expressed the MAMP reporter (as red fluorescent protein dTomato) and FER-GFP. We found that almost all MAMP-responsive cells accumulated FER-GFP in the nucleus after CHA0 and DC3000 treatment (Fig. 1h-i). An immunoblot analysis detected FER C-terminal region containing the intracellular kinase domain after CHA0 and DC3000 treatment (Extended Data Fig. 1f), which was named FERN (FER in the Nucleus). Together, these results indicate that FER relocation from PM to the nucleus is linked to LI to modulate rhizobacteria colonization in the root.

### Rhizobacteria trigger RALF23 accumulation and FER cleavage

Given that bacteria can trigger FER cleavage and that RALFs serve as FER’s ligand ^17,31^, we investigated whether bacterial-induced FER cleavage is somehow linked to levels of RALFs. Ten *RALF* genes, including *RALF23*, were expressed in Arabidopsis roots (Extended Data Fig. 2a) ^31^. We used RALF23, which was shown to regulate immunity in the leaves ^16,17^, as a model peptide to analyze its levels in response to bacteria colonization. Treatment of DC3000 or CHA0 induced the accumulation of mature RALF23 in *pRALF23::RALF23-GFP* seedlings (Fig. 2a), especially in the TZ and EZ of roots (Extended Data Fig. 2b). The function of RALF23 requires proteolytic cleavage to release its C-terminal mature peptide ^32^. To estimate mature RALF23-GFP absolute abundance, recombinant S-tag-GFP (S-GFP) was expressed in *E. coli* and purified. We found that DC3000 and CHA0 induced an increase in mature RALF23-GFP accumulation from 10-50 nM under rest conditions to a range of 80-200 nM (Extended Data Fig. 2c). To examine whether elevated level of the mature RALFs was responsible for FER nuclear localization and LI, we employed an *s1p* mutant in which cleavage of RALF23 was compromised (Extended Data Fig. 2d) ^17,32^ to study the relationship of RALF23 maturation and production of FERN in *s1p* mutant. We found that FERN induced by DC3000 and CHA0 in *s1p* mutant is significantly reduced relative to WT (Extended Data Fig. 2e). Interestingly, exogenous application of synthetic RALF23 at a concentration of 200 nM restored the cleavage activity of FER in *s1p* mutant in response to DC3000 and CHA0 (Extended Data Fig. 2e), suggesting that CHA0 or DC3000-induced FER nuclear localization might be attributed to the enhanced maturation of the RALF ligand (e.g., RALF23).

**Fig. 2.**
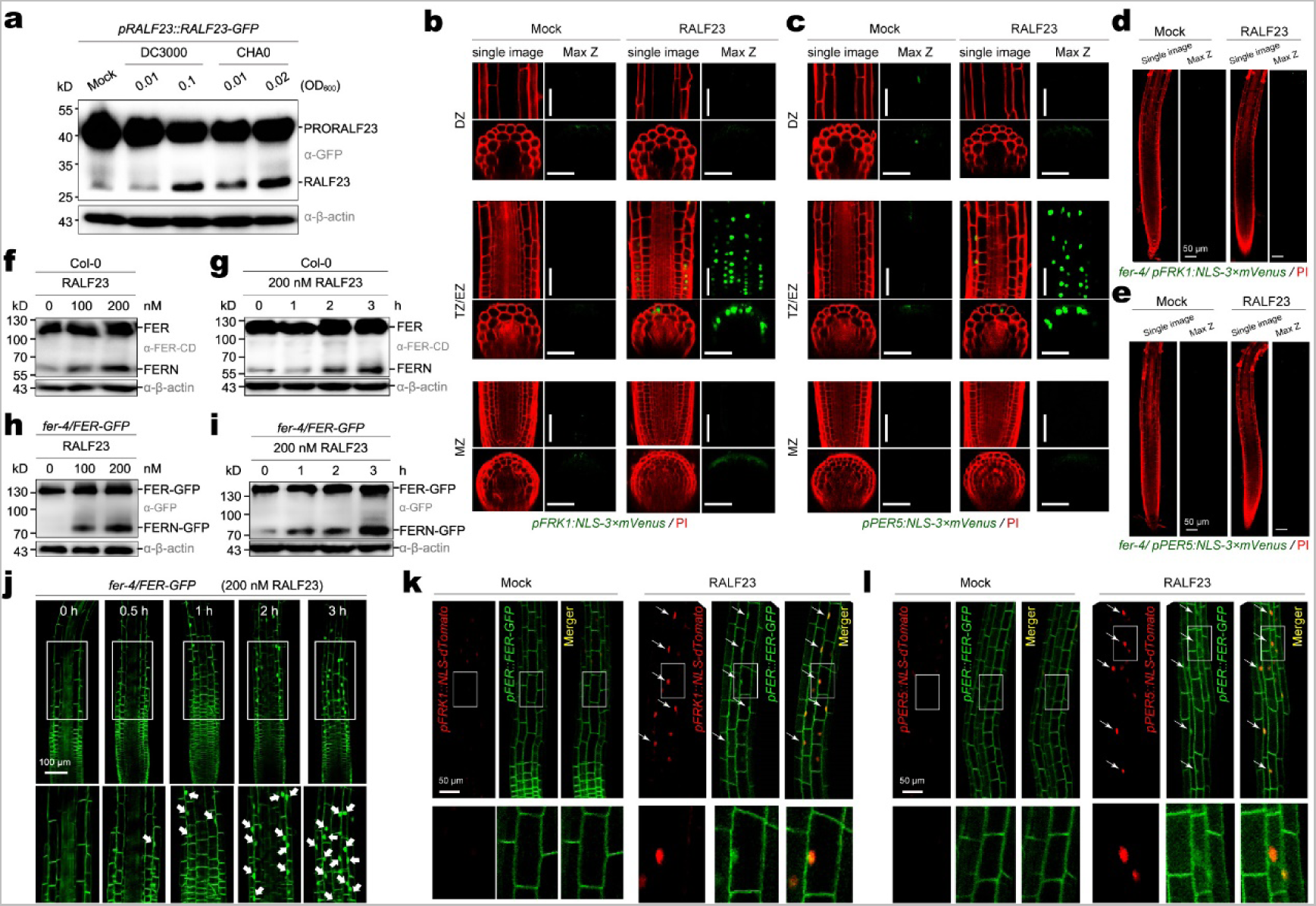
RALF23 induces the cleavage and nuclear localization of FER intracellular domain in the TZ and EZ. (**a**) Immunoblot analysis of RALF23 cleavage in *pRALF23::RALF23-GFP* seedlings in response to DC3000 or CHA0 for 8 h in D-roots. (**b–c**) mVenus fluorescence patterns in *pFRK1::NLS-3×mVenus* (**b**) or *pPER5::NLS-3×mVenus* (**c**) lines in response to 200 nM RALF23 for 12 h in the Col-0 background. mVenus signals co-visualized with PI. Single confocal microscope image and maximal projections of Z-stacks are presented. Scale bar, 50 μm. (**d–e**) mVenus fluorescence patterns in *pFRK1::NLS-3×mVenus* (**d**) or *pPER5::NLS-3×mVenus* (**e**) lines in response to 0.2 μM RALF23 for 12 h in the *fer-4* mutant background. mVenus signals co-visualized with PI. (**f–g**) FER cleavage in Col-0 (**f**) and *fer-4/FER-GFP* (**g**) seedlings in response to RALF23 treatment for 3 h in D-roots. The C-terminal bands were detected using α-FER-CD antibody (**f**), or α-GFP antibody (g). (**h–i**) Col-0 **(h)** and *fer-4/FER-GFP* **(i)** seedlings treated with RALF23 along a time course. (**j**) GFP fluorescence localization in the D-roots of *fer-4/FER-GFP* treated with 200 nM RALF23 along a time course. (**k–l**) FERN co-localizes with FRK1 (**k**) or PER5 (**l**) in the TZ and EZ cells after treatment with 200 nM RALF23 for 12 h. Arrows indicate the nucleus.

To further confirm the induction of FER cleavage by RALF23 is associated with LI, we initially validated that flg22 also induced the accumulation of RALF23 to a concentration range of 80-200 nM from 10-50 nM under rest conditions (Extended Data Fig. 2f-h) ^17^. Next, we examined whether RALF23 influences flg22-triggered immunity by applying varying levels of exogenous RALF23. Intriguingly, RALF23 in the range of 100-200 nM promoted the expression of immunity reporter genes induced by flg22 (Extended Data Fig. 3a-b), whereas increased concentrations of RALF23 (1 μM or higher) significantly inhibited immunity (Extended Data Fig. 3c-d) ^17^. Treatment with RALF23 (200 nM) primarily triggered the expression of reporter genes in TZ and EZ of roots (Fig. 2b-c), indicating that RALF23 induces root immune responses in specific regions. We further showed that the RALF23-induced LI was absent in *fer-4* mutant (Fig. 2d-e). Interestingly, using *fer-4/FER-GFP* plants, we found that 100-200 nM RALF23 triggered nuclear accumulation of FER C-terminal product FERN (Fig. 2f-i). FERN again was undetectable when the RALF23 concentration exceeded the endogenous physiological level (>500 nM, Extended Data Fig. 3e). Similar to CHA0 and DC3000 treatments, RALF23 treatment resulted in clearly detectable fluorescence of FERN-GFP in the nucleus (Fig. 2j, Extended Data Fig. 3f). The fluorescence FERN-GFP co-localized with MAMP reporter-dTomato (Fig. 2k-l). In parallel, we detected more FERN in the nuclear fraction relative to the cytoplasmic fraction after 200 nM RALF23 treatment (Extended Data Fig. 3g). Previous research has shown that higher RALF23 concentration (1000 nM) suppresses immunity ^17,33^.

To confirm concentration-dependent effects of RALF23 on root immune responses, we analyzed ROS levels and FLS2-BAK1 interaction in response to different concentrations of RALF23. We found that lower doses of RALF23 (100-200 nM) boosted ROS levels, whereas higher levels (500-1000 nM) inhibited ROS production (Extended Data Fig. 4a). We further showed that RALF23-mediated regulation of ROS production depends on FER (Extended Data Fig. 4b). Additionally, lower RALF23 concentrations (100-200 nM) enhanced the FLS2-BAK1 interaction triggered by flg22, whereas higher concentrations (500-1000 nM) hindered this interaction (Extended Data Fig. 4c). This suggested that RALF23 at different concentrations has opposite effects on root-localized immunity, consistent with previous findings ^17^. We also observed that under low-phosphate (LP) conditions, FER cleavage in response to both CHA0 and DC3000 treatments was significantly reduced compared to the normal condition, which aligned with previous work showing that LP inhibits root immunity via RALF23-FER ^34^(Extended Data Fig. 4d). Under LP and normal conditions, both CHA0 and DC3000 can induce RALF23 maturation (Extended Data Fig. 4e). However, in LP conditions, the degree of RALF23 maturation is comparatively lower than under normal conditions (Extended Data Fig. 4e). Taken together, these results suggest that FER may suppress immunity when RALF23 level is low, and promote immunity when RALF23 level increases to induce cleavage and relocation of FER C-terminal domain into the nucleus in TZ and EZ cells. In summary, DC3000 and CHA0 enhanced RALF23 maturation, leading to FER cleavage.

### Zinc metalloproteinase At2-MMP cleaves FER

To dissect the mechanism of FER cleavage, we screened the candidate proteases that may use FER as a substrate. We developed a protease cleavage screening system by monitoring FER-GFP fluorescence in the nucleus. The Arabidopsis genome encodes approximately 826 proteases ^35^. Screening for protease genes expressed in roots, we identified a zinc metalloproteinase At2-MMP (At1g70170) to be essential for nuclear FER-GFP signal (Extended Data Fig. 5a-b, Extended Data Fig. 6a). Moreover, BB-94, a broad-spectrum MMP inhibitor ^36^, blocked RALF23-triggered FERN production (Extended Data Fig. 6b), which supported the role of At2-MMP in FER cleavage.

The conserved zinc binding motif HEXXHXXGXXH within the catalytic domain is required for MMP activity ^37^. We produced and purified the recombinant At2-MMP catalytic domain (At2-MMP-CD) and the inactive variant of At2-MMP-CD protein (At2-MMP-CD^mut^). The recombinant FER proteins were incubated together with either wild type or inactive variant of At2-MMP-CD (Extended Data Fig. 6c-d) to perform FER cleavage assay. While the At2-MMP-CD wild type produced more cleavage product of FER_225–808_ along the time course (Fig. 3a), the dead variant At2-MMP-CD failed to cleave FER_225–490_ or FER_521–895_ (Extended Data Fig. 6e). We then synthesized a fragment of FER (487-526, termed as FER intracellular juxtamembrane [FER-IJM]) and identified three cleavage sites (S493/L494, A513/S514, and P517/S518; Fig. 3b, Extended Data Fig. 6f). The FER variant lacking the cleavage site (FER^del3^_225–808_) was not cleaved by At2-MMP-CD *in vitro* (Fig. 3c, Extended Data Fig. 6e). Subsequent investigations revealed that under normal conditions, At2-MMP is not expressed in TZ and EZ (Extended Data Fig. 7a). However, following treatments with DC3000 and CHA0, a significant upregulation of At2-MMP within the TZ and EZ regions was observed, primarily within the epidermis of *pAt2-MMP::NLS-dTomato* seedlings (Extended Data Fig. 7a). This observation suggests that the induction of FER cleavage within TZ and EZ by DC3000 or CHA0 may indeed be attributed to At2-MMP. Further validation was performed by analyzing two mutant lines (*at2-mmp-1* and *at2-mmp-2*) along with two transgenic lines overexpressing *At2-MMP* (*At2-MMP-OE#2* and *At2-MMP-OE#3*; Extended Data Fig. 7b-c). These experiments confirmed a direct correlation between At2-MMP levels, the cleavage of FER upon treatment with CHA0 and DC3000 (Fig. 3d), and the LI responses (Fig. 3e). Additionally, treatment with RALF23 also yielded results consistent with the responses observed following CHA0 and DC3000 treatment (Extended Data Fig. 7d-f). It is worth noting that the cleavage sites in the IJM domain are conserved across different plant species (Extended Data Fig. 8a). In tobacco (*Nicotiana tabacum*), a paralog of FER known as NtFER (LOC107813068) also exhibited cleavage upon treatment with RALF23 (Extended Data Fig. 8b). The activity of MMPs relies on cleavage of the cysteine switch ^38,39^. Unlike full-length At2-MMP-4×myc, truncated At2-MMP accumulated after treatment with RALF23 (Extended Data Fig. 9a). Moreover, total protein extracted from *At2-MMP-*OE seedlings treated with RALF23 catalyzed more cleavage of recombinant glutathione S-transferase (GST)-FER_225-808_ *in vitro* (Extended Data Fig. 9b). Collectively, these findings support the notion that CHA0 and DC3000 may induce and activate At2-MMP in the TZ and EZ, subsequently leading to FER cleavage.

**Fig. 3.**
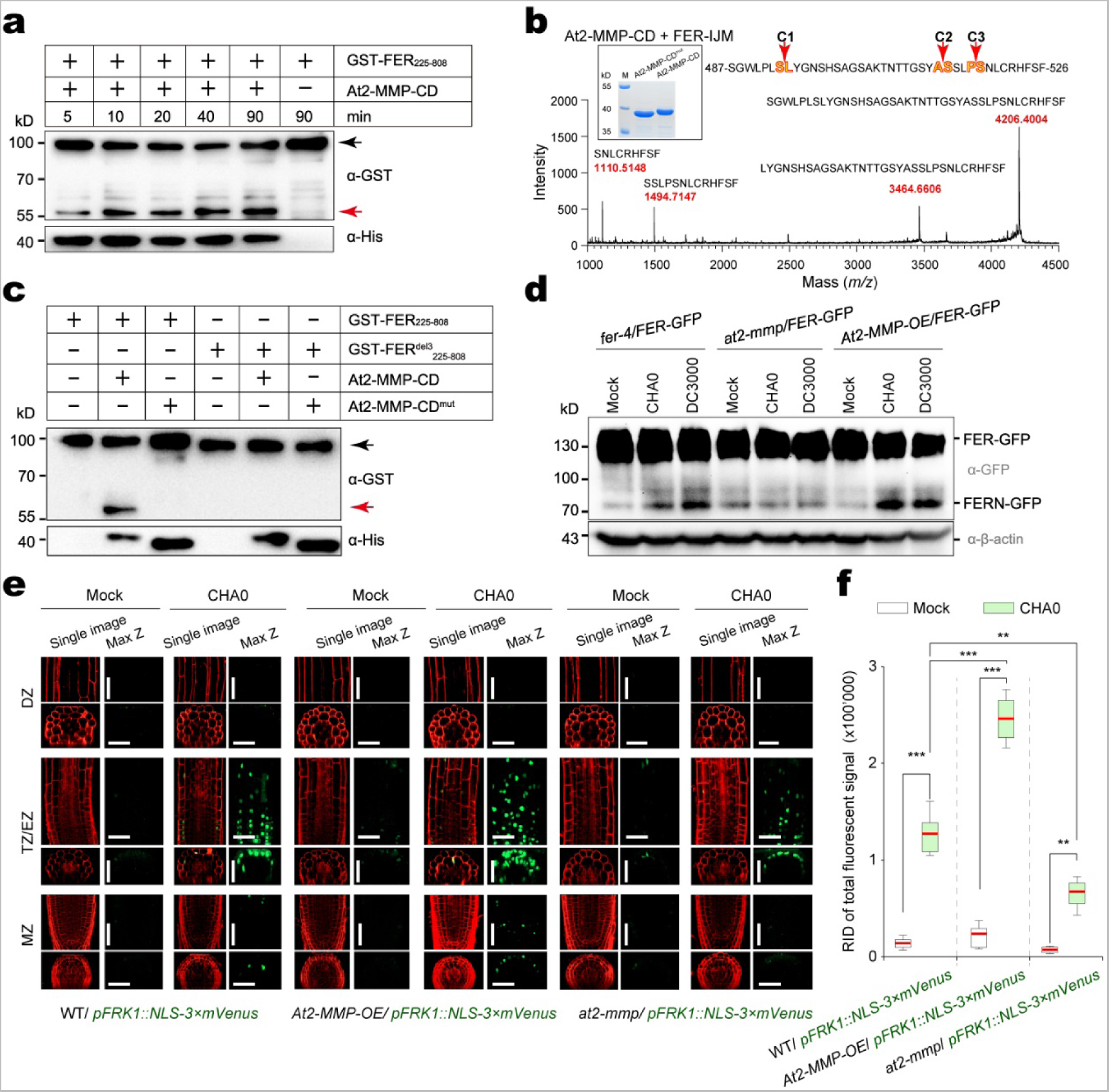
At2-MMP cleaves FER, mediating CHA0 and DC3000-driven localized immune responses. (**a**) Proteolytic activity of recombinant At2-MMP-CD toward FER_225–808_. The red arrow represents the cleaved N-terminal of GST-FER, and the black arrow represents the full-length GST-FER. (**b**) MALDI-TOF MS analysis of FER-IJM cleavage by At2-MMP-CD *in vitro*. Amino acid sequences and corresponding molecular weights of FER-IJM and FER-IJM peptides cleaved by At2-MMP-CD are indicated. The insert shows the SDS-PAGE analysis of purified At2-MMP-CD and At2-MMP-CD^mut^. (**c**) Proteolytic activity of recombinant His-At2-MMP and His-At2-MMP^mut^ toward GST-FER_225–808_ and GST-FER^del3^_225–808_. (**d**) At2-MMP cleaves FER in plants. *fer-4/FER-GFP*, *at2-mmp-1* and *At2-MMP-*OE #2 in the *fer-4/FER-GFP* background in response to DC3000 or CHA0 for 8 h in D-roots. (**e**) mVenus fluorescence pattern of the MAMP promoter marker line *pFRK1::NLS-3×mVenus* in response to DC3000 or CHA0 for 18 h in the Col-0, *At2-MMP-*OE #2, and *at2-mmp-1* backgrounds. mVenus signals co-visualized with PI. Scale bar, 50 μm. (**f**) Quantitative analysis of mVenus signal intensities of the *FRK1* marker in response to DC3000 or CHA0 for 18 h at TZ/EZ in the Col-0, *At2-MMP-*OE #2, and *at2-mmp-1* backgrounds. Boxplot centers show median (n = 12 roots). Asterisks (***p < 0.001, **p < 0.01) indicate statistically significant differences between means by ANOVA and Tukey’s test analysis.

### FERN-dependent LI inhibits bacterial colonization

To dissect the role(s) of nucleus-localized FERN in growth and development, we generated transgenic lines harboring several variants of FER in the *fer-4* background (Extended Data Fig. 10a-b). The FERN-GFP protein was localized to both cytosol and nucleus (Extended Data Fig. 10b), whereas FERN-GFP-NES (with a nucleus export signal) accumulated mainly in the cytosol (Extended Data Fig. 10b). Expression of FERN-GFP or FERN-GFP-NES failed to complement the defective phenotypes of rosette and root hair in *fer-4* (Fig. 4a and d). But FERN-GFP restored root morphology in the MZ and TZ of the *fer-4* mutant (Fig. 4e-f). The FER^ΔJM^-GFP variant featured a deletion of the IJM domain, preventing FER cleavage in response to RALF23 (Extended Data Fig. 10c). Hence, FER^ΔJM^-GFP exhibited continuous localization at the PM (Extended Data Fig. 10d), and this variant restored root hair, overall growth, and reproduction phenotypes in *fer-4* (Fig. 4 a-d). Moreover, it also reinstated the root morphology in the MZ and TZ of *fer-4* (Fig. 4e-f). In summary, PM-FER facilitates plant cell growth and suppresses immunity, while FERN primarily activates LI and has minimal impact on overall cell growth beyond the TZ and EZ of roots.

**Fig. 4.**
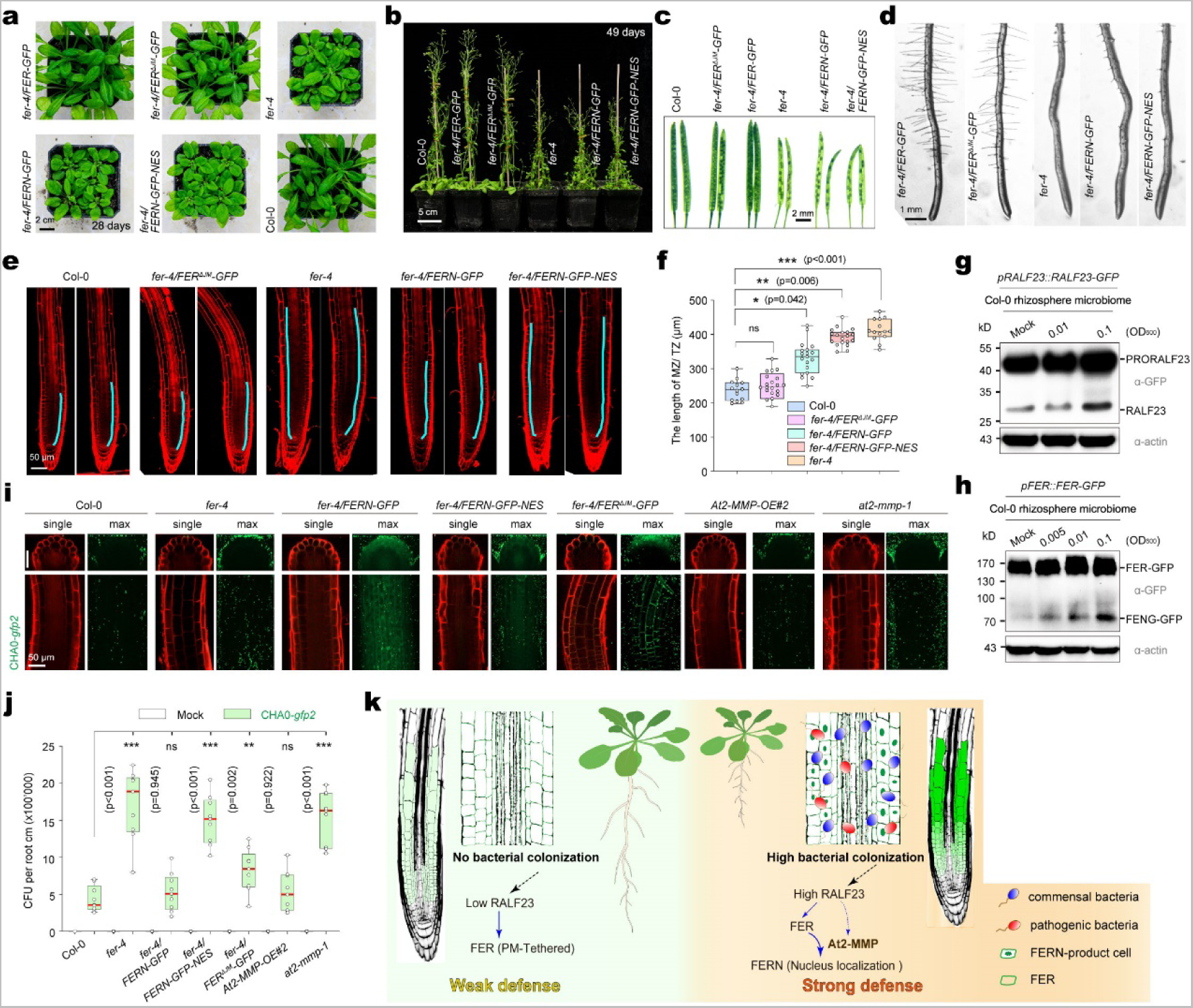
FERN regulates localized immune responses during root-bacteria communication. (**a**) Representative images of Col-0, *fer-4*, *fer-4/FER-GFP*, *fer-4/FER^ΔJM^-GFP*, *fer-4/FERN-GFP*, *fer-4/FERN-GFP-NES* plants grown on soil for 28 days under white light (16-h light/8-h dark cycles). (**b**) Representative images of Col-0, *fer-4*, *fer-4/FER-GFP*, *fer-4/FER^ΔJM^-GFP*, *fer-4/FERN-GFP*, *fer-4/FERN-GFP-NES* plants grown on soil for 49 days under white light. (**c**) Representative images of Col-0, *fer-4*, *fer-4/FER-GFP*, *fer-4/FER^ΔJM^-GFP*, *fer-4/FERN-GFP*, *fer-4/FERN-GFP-NES* siliques. (**d**) Root hairs from 5-day-old of *fer-4*, *fer-4/FER-GFP*, *fer-4/FER^ΔJM^-GFP*, *fer-4/FERN-GFP*, *fer-4/FERN-GFP-NES* lines seedlings. (**e**) Primary roots of Col-0, *fer-4*, *fer-4/FERN-GFP*, *fer-4/FERN-GFP-NES*, and *fer-4/FER^ΔJM^-GFP* lines in 7-day-old D-seedlings. The thick cyan lines represent MZ and TZ cells in the root cortex. Root cells are highlighted by PI staining. (**f**) Cell length in the MZ and TZ of 7-day-old roots in (**e**). Values are means ± SD, *n* ≥ 14, two-tail Student’s t-test (\**p* < 0.05, \*\**p* < 0.01, and \*\*\**p* < 0.001), ns, not significant. (**g**) Immunoblot analysis of RALF23 cleavage in *pRALF23::RALF23-GFP* seedlings in response to *fer-4* rhizosphere microbiome or Col-0 rhizosphere microbiome for 8 h in D-roots. (**h**) Immunoblot analysis of RALF23 cleavage in *pFER::FER-GFP* seedlings in response to *fer-4* rhizosphere microbiome or Col-0 rhizosphere microbiome for 8 h in D-roots. (**i**) Representative images showing CHA0-*gfp2* colonization on Col-0, *fer-4*, *fer-4/FER^ΔJM^-GFP*, *fer-4/FERN-GFP*, *fer-4/FERN-GFP-NES*, *at2-mmp-1*, and *At2-MMP-*OE#2 roots in the TZ and EZ at 3 dpi. Photographs are maximum projections of confocal Z stacks. (**j**) Colony forming units (CFU) counting of CHA0-*gfp2* colonization on Col-0, *fer-4*, *fer-4/FER^ΔJM^-GFP*, *fer-4/FERN-GFP*, *fer-4/FERN-GFP-NES*, *at2-mmp-1*, and *At2-MMP-*OE#2 roots in the TZ and EZ at 3 dpi. Three roots were collected for each sample at the indicated colonization time point. Values are means ± SD (3 biological replicates). Asterisks (\*\**p* < 0.01, and \*\*\**p* < 0.001) indicate significant differences based on ANOVA followed by Tukey’s test. ns, not significant. (**k**) Diagram showing that a high concentration of mature RALF23 induces FER cleavage via At2-MMP to produce FERN, which relocates to the nucleus and activates immune responses in the TZ and EZ.

To investigate the effects of WT rhizosphere microbiome (from the rhizosphere of 4-week-old WT plants by shaking the soil in sterile water) on the maturation of RALF23 and the cleavage of FER, we treated *pRALF23::RALF23-GFP* and *pFER::FER-GFP* seedlings with separated Col-0 rhizosphere microbiome. The results indicated that the Col-0 microbiome significantly enhanced RALF23 maturation and FER cleavage (Fig. 4 g-h). We next tested whether FERN nuclear localization affected bacterial colonization and found that CHA0 grew more in the TZ and EZ of *fer-4*, *fer-4/FER^ΔJM^-GFP*, *fer-4/FERN-GFP-NES*, and *at2-mmp*, but less in Col-0, *fer-4/FERN-GFP* and *At2-MMP-*OE (Fig. 4 i-j), indicating that FERN inhibits bacterial colonization around TZ and EZ. These results further supported the conclusion that PM-localized FER and nucleus-localized FERN showed distinct effect on colonization of CHA0 around TZ and EZ, an indicator of LI.

The *fer-4* mutant is relatively specifically enriched for pseudomonas fluorescens and forms a bacterial community that helps promote the growth of the next generation of Arabidopsis ^33^. To address whether the presence of FERN alters the rhizosphere microbiome associated with the plants in natural soil, we performed microbiome transplant assays ^34^ and observed a growth promotion effect on wild-type plants grown in the presence of the microbiome associated with the seedlings of *fer-4*, *fer-4/FERN-GFP-NES*, *fer-4/FER^ΔJM^-GFP*, and *at2-mmp-1*, but not on Col-0, *fer-4/FERN-GFP*, or *At2-MMP-*OE#2 (Extended Data Fig. 10e), further confirming the roles of FERN in rhizosphere bacterial colonization of roots.

## Discussion

We have identified a new immune signaling pathway in which the receptor kinase FER controls LI. A working model was proposed (Fig. 4k): In the absence of microbial signals, mature RALF23 is kept at a low level and full-length FER is present in the PM to maintain normal plant growth. Upon detection of high bacterial colonization or external flg22 application, mature RALF23 accumulates and triggers the cleavage and nuclear accumulation of FERN that, in turn, enhances the expression of immunity-related genes in a restricted region of roots, thus protecting these vulnerable areas from microbe invasion. Our results provide insights into a mechanism by which plants interact with and optimize bacterial communities in roots through the opposite roles of FER and FERN in immunity.

In our study, we found that different concentrations of RALF23 have distinct effects on the immune response. RALF23 concentrations at 100-200 nM promote ROS production and enhance FLS2-BAK1 interaction, indicating a positive role in immune regulation. In contrast, higher RALF23 concentrations at 500-1000 nM inhibit ROS generation and weaken FLS2-BAK1 interaction (Extended Data Fig. 4a-c), consistent with previous findings^17,33^. Moreover, under low-phosphate conditions, CHA0 or DC3000 also induces RALF23 maturation but fails to stimulate FER cleavage, thus impeding LI (Extended Data Fig. 4d-e). Our research introduced the intriguing concept that bacterial colonization plays a critical role in determining RALF levels by an unknown mechanism, offering a critical area for future research in plant-microbe interactions. Our analysis suggests that localized expression of RALF23 and At2-MMP could be a prominent factor contributing to the observed localized immunity in the TZ/EZ of roots (Extended Data Fig. 2b; Extended Data Fig. 7a), but comprehensive mechanistic studies are warranted to link these events and FER cleavage to the downstream immunity programs in plant cells.

## Acknowledgments

We thank Dr. Alice Cheung, Dr. Gregor Langen and Dr. Michiael R. Sussman for providing plant materials;

## Authors’ contributions

F. Y. and J. C. conceived the project and designed research; J. C., X-N. Q., H-B. L., F. X., L. W., L-L. J., C-Y. L., B-Q. W. and D-S. W performed research; S. L and F. Z contributed new ideas; J. C. and F. Y. analyzed data and wrote the paper; all authors reviewed and approved the manuscript for publication.

## Data and materials availability

All data are available in the manuscript or the supplementary materials.

## Materials and Methods

### Plant materials and growth conditions

All seeds were surface sterilized and stratified at 4°C for 3 days and then grown on half-strength Murashige and Skoog (MS) medium with 0.8% (w/v) sucrose solidified with 1% (w/v) agar (A7002, Sigma-Aldrich) for analysis. Arabidopsis (*Arabidopsis thaliana*) wild-type (Col-0) was the accession used in this study. The *fer-4* mutant ^23^ and the *pFER::FER-GFP* line in the *fer-4* background ^23^ were reported previously. The *pRALF23::RALF23-GFP* constructs were cloned into the plant expression vector p1300-GFP with *EcoR*I and *BamH*I. The *at2-mmp-1* (GK-416E03.01) and *at2-mmp-2* (SALK_082450) mutants were obtained from the ABRC and were confirmed by PCR with specific primers (Table S1). The *pFRK1::NLS-3×mVenus* and *pPER5::NLS-3×mVenus* lines were obtained from Dr. Niko Geldner’s Lab and were crossed to *fer-4*, *At2-MMP-*OE#2 and *at2-mmp-1*. The *FERN-GFP* and *FERN-GFP-NES* constructs were cloned into the plant expression vector pDT7 ^40^ with *Spe*I and *Pac*I, the *At2-MMP* coding sequence was cloned into pDT7 with *Spe*I to generate *ACTIN2* promoter-driven constructs. The nuclear export signal (NES) sequence was generated by PCR to encode the amino acid residues LQNELALKLAGLDINKTGG (stop) ^41^. *pACTIN2::FERN-GFP* and *pACTIN2::FERN-GFP-NES* plants were crossed to *fer-4*. The *pFER::FER^ΔJM^-GFP* construct was generated in the plant expression vector p1300-GFP with *EcoR*I and *BamH*I. The *pFRK1::NLS-dTomato* and *pPER5::NLS-dTomato* constructs were generated into the plant expression vector pCAMBIA1300 with *EcoR*I and *Sal*I to generate the *FRK1* and *PER5* promoter-driven plants, which were then crossed into the *fer-4/FER-GFP* background. Positive transformants were selected by resistance to basta or kanamycin and verified by immunoblotting using α-GFP (Abclonal, AE012) or α-Myc (Cell Signaling Technology, 2040) antibodies. All primers used for plasmid constructions and transgenic lines are listed in Table S1.

Seedlings for root growth inhibition or MAMP marker lines assays were first grown vertically on half-strength MS medium solidified with 1% (w/v) agar at 22°C under 16-h light/8-h dark cycles. The square plate was introduced into the methacrylate box and the methacrylate comb was inserted into the agar to block the light from the top fluorescents for 4 days (dark root, D-root) ^42^, then seedlings with similar root size were transferred to liquid half-strength MS medium containing the mentioned peptide molecules or bacteria using 24-well culture plates.

### Microscopy settings and image processing

Confocal laser scanning microscopy was performed on a Zeiss LSM880 inverted confocal scanning microscope. DAPI; mVenus/GFP; PI and dTomato were excited at 350 nm, 488 nm and 543 nm wavelength, respectively. Photographs were taken with 20× or 40× water immersion objective. For more detailed analyses in large areas of interest, imaging was performed using Z-scan with tile-scan (overlap 10%). Sequential scanning was used to avoid interference between fluorescence channels. Confocal images after treatments were taken following identical criteria.

For microscopy analysis of MAMP marker lines under various treatments, 5-day-old seedlings grown in the D-root device were carefully transferred to liquid half-strength MS medium containing peptides or bacterial strains using 24-well culture plates (the surface of the 24-well plate was covered with tin foil, a 0.5-mm diameter hole was punched through the foil; seedling roots were carefully moved into the hole to ensure D-root). The seedlings were observed under confocal microscopy 18 h after treatment. A pool of 10-12 homozygous seedlings were analyzed for each assay. At least three independent replicates were performed.

### RNA extraction, cDNA synthesis and quantitative PCR

Root samples from the different treatments were ground in liquid nitrogen for RNA extraction. Total RNA was extracted using TRIzol reagent (Ambion, 15596-026) according to the manufacturer’s instructions and treated with DNase I (Takara) to remove genomic DNA. RNA samples were quantified with a Nanodrop spectrophotometer (Thermo Scientific). First-strand cDNA was synthesized from 3 μg total RNA using a Maxima H Minus First Strand cDNA Synthesis Kit with dsDNase (Thermo Scientific, K1682) and an oligo(dT) primer, according to the manufacturer’s instructions. cDNA was amplified in triplicate by quantitative PCR using a 2× AceQ SYBRGreen qRCR Premix (with ROX) (Vazyme). *ACTIN2* was used as a reference gene in qPCR analysis. Primers used for qPCR are listed in Table S1.

### RALFs and bacterial infection-induced FER cleavage analyses

RALF1/23 were synthesized by Guoping Pharmaceutical Co., LTD and dissolved in water. For RALFs peptide-induced cleavage of FER, 5-day-old seedlings grown in the D-root system were treated in liquid half-strength MS medium containing different RALF peptides for 3 h at 22°C in the dark. Then, roots from all seedlings were collected and incubated in extraction buffer (20 mM HEPES pH 7.5, 150 mM NaCl, 10% [v/v] glycerol, 1% [v/v] Triton X-100, 5 mM EDTA, 10 mM DTT, 1 mM PMSF and protease inhibitor cocktail [Halt Protease Inhibitor Cocktail, 78430, Thermo Fisher Scientific Inc.]) for 1 h. Then, samples were centrifuged at 12,000 rpm for 10 min at 4°C, supernatants were collected, and 4× SDS loading buffer was added and samples were boiled for 10 min at 95°C. Analysis was carried out by 10% SDS-PAGE and immunoblotting using α-FERCD (6×His-FER-CD was purified and used as an antigen to produce a polyclonal antibody in mouse), α-GFP or α-ACTIN (plant-specific) (Abclonal, AC009) antibodies.

For the effect of bacterial infection on FER cleavage, uniform 5-day-old *fer-4/FER-GFP* seedlings were selected and transferred to bacterial suspension at density of OD_600_ =0.01 or 0.1 for DC3000-*gfp2*, OD_600_ = 0.01 or 0.02 for CHA0-*gfp2* for 8 h. After infection, the bacterial solution was carefully removed from the roots by washing with sterile MiliQ water before immunoblotting or confocal imaging.

### Quantitative determination of mature RALF23 in *pRALF23:RALF23-GFP* seedlings

Mature RALF23 was quantified as described ^43^ with a few modifications. *pRALF23:RALF23-GFP* seedlings were grown vertically on half-strength MS medium solidified with 1% (w/v) agar at 22°C under a 16-h light/8-h dark cycles for 5 days in the D-root device. Then, 0.15 g of seedlings (about 120 μL total volume) was collected, gently ground, and centrifuged at 12,000 rpm for 10 min. The supernatants were collected and incubated in 120 μL extraction buffer (20 mM HEPES pH 7.5, 150 mM NaCl, 10% [v/v] glycerol, 1% [v/v] Triton X-100, 5 mM EDTA, 10 mM DTT, 1 mM PMSF and protease inhibitor cocktail) for 1 h. Then, the samples were centrifuged at 12,000 rpm for 10 min, supernatants were collected, and 4× SDS loading buffer was added (40 μL) and the samples were boiled for 10 min at 95°C. Analysis was carried out by 11% SDS-PAGE and immunoblotting using an α-GFP antibody. The S-GFP standard was cloned into pET-32a using *BamH*I and *EcoR*I restriction enzymes and purified using Ni NTA Beads (Smart Life Sciences, SA004100); protein concentration was determined by the BCA method, and the gray value of mature RALF23 was converted into concentration through S-GFP gray value from immunoblots. The concentration of mature RALF23 in *pRALF23:RALF23-GFP* seedlings was obtained by the ratio of the total volume of extracted roots to the volume of loaded samples. The concentration of endogenous mature RALF23 peptide was in the nanomolar range under both resting (10-50 nM) and flg22, CHA0 or DC3000 treatments (80-200 nM) in D-roots.

Fluorescence intensity was used to measure the concentration of mature RALF23 in *pRALF23:RALF23-GFP* seedlings, using the ratio of the gray value of full-length RALF23 and mature RALF23 in *pRALF23:RALF23-GFP* seedlings. The S-GFP standard was selected as the standard concentration. The storage concentration was 100 μM, and a confocal microscope was used to prepare the standard curve of GFP protein. To adapt to the fluorescence intensity of RALF23-GFP and avoid oversaturation, the maximum molar concentration of the GFP standard curve was controlled at approximately 2.5 μM. The concentration of mature RALF23 in *pRALF23:RALF23-GFP* seedlings was obtained according to the GFP standard curve.

### Fractionation of subcellular organelles

Nuclear and cytosolic fractionation was performed as described previously with minor modifications ^44^. 5-day-old D-root seedlings were transferred to liquid half-strength MS medium with RALF peptides for 3 h at 22°C in the dark. Then, roots from all seedlings were collected and extracted with extraction buffer (20 mM Tris-HCl pH 7.0, 250 mM sucrose, 25% [v/v] glycerol, 20 mM KCl, 2 mM EDTA, 2.5 mM MgCl_2_, 1 mM DTT, 1× protease inhibitor cocktail and 1% [v/v] Triton X-100). The obtained solution was centrifuged at 3,000 g for 5 min. The resulting supernatant was used as the cytosolic fraction and the pellet was washed with resuspension buffer (20 mM Tris-HCl pH 7.0, 25% [v/v] glycerol, 2.5 mM MgCl_2_ and 1 mM DTT) to obtain the nuclear fraction. Protein samples were separated on 12% SDS-PAGE and probed with α-GFP, α-GAPDH (ABclonal, AC002) (Marker antibody of cytoplasm), or α-histone H3 (Abcam, ab1791) (Marker antibody of the nucleus) antibodies with a corresponding dilution ratio at 4°C overnight. The membrane was incubated with goat α-rabbit (Jackson Immuno Research, 111-035-003) or α-mouse (Jackson Immuno Research, 115-035-003) HRP-conjugated secondary antibodies against the primary antiserum for 1 h.

### Protoplast transient expression assay

Protoplasts were prepared from 7-day-old Col-0 roots. About 40 μg of DNA plasmids, consisting of pairs of the *pFER::FER-GFP* constructs plus a single protease gene, were individually transfected into protoplasts. Protease genes that were predicted to be expressed in Arabidopsis roots were selected, resulting in a list of 203 protease genes. The transfected protoplasts were incubated in the dark at 23°C for 18 h to allow protein production. Subsequent analyses were conducted using a laser-scanning confocal microscope (Zeiss 880).

### *In vitro* cleavage assay

For the *in vitro* cleavage assays, FER_225–808_, GST-FER_225–490_, GST-FER_225–536_, GST-FER_521–895_, GST-FER^del1^_225–808_, GST-FER^del2^_225–808_ and GST-FER^del3^_225–808_ were recombined into the pGEX-4T-1 vector; the At2-MMP catalytic domain corresponding to the predicted mature At2-MMP (At2-MMP-CD) was recombined into the pET-32a vector. The mutated version of the catalytic domain of At2-MMP (i.e., zinc-binding sequence <u>HE</u>XX<u>H</u>XX<u>G</u>XX<u>H</u> with the H/E/H/G/H residues replaced with Ala [A]) was recombined into the pET-32a vector to produce At2-MMP-CD^mut^. All recombinant proteins were produced in *Escherichia coli* and purified with glutathione beads (Smart Life Sciences, SA008100) or Ni NTA Beads affinity resins. For *in vitro* cleavage assays, 10 μg of GST-FER_225–808_, GST-FER_225–490_, GST-FER_225–536_, GST-FER_521–895_, GST-FER^del1^_225–808_, GST-FER^del2^_225–808_, or GST-FER^del3^_225–808_ and 10 μg of At2-MMP-CD or At2-MMP-CD^mut^ recombinant proteins were incubated in a tube at 38℃ for the indicated time in a 50 μL reaction. The concentration of the indicated chemicals was as follows: 50 mM Tris-HCl (pH 7.5), 10 mM CaCl_2_, 1 mM EDTA and 50 μM ZnCl_2_. Protease activity was inhibited by adding 50 mM EDTA. Proteins and their cleavage products were detected by Coomassie brilliant blue staining and immunoblotting using α-GST antibody (ABclonal, AE001).

### Determination of cleavage sites by mass spectrometry

Cleavage sites of the peptide fragment of FER from Ser-487 (S487) to Phe-526 (F526) (Named FER-IJM) was determined by MALDI-TOF MS (AB SCIEX TOF/TOF TM 5800 system, Applied Biosystems, USA). First, 10 μg of FER-IJM and 10 μg of At2-MMP-CD or At2-MMP-CD^mut^ recombinant proteins were incubated at 38℃ for 30 min in a 50 μL reaction. A 1 μL aliquot of each reaction was spotted onto a 384-well target plate along with an equal volume of a matrix solution containing 10 mg/mL a-cyano-4-hydroxycinnamic acid, 50% (v/v) acetonitrile and 0.1% (w/v) trifluoroacetic acid (TFA). The mixture was allowed to dry at room temperature. Calibration of the instrument was performed externally with a Peptide Mass Standard Kit. The monoisotopic molecular mass of each peptide was determined.

### Bacterial strains and growth conditions

*Pseudomonas protegens* strain CHA0 is a tobacco (*Nicotiana tabacum*) root isolate with plant-beneficial activities. The bacterial pathogen *Pseudomonas syringae* pv. tomato DC3000 is a phytobacterial pathogen. Bacterial strains were incubated overnight in liquid LB medium at 28℃. Bacterial cells were collected by centrifugation and resuspended in sterile MiliQ water for root inoculation assays.

The GFP-tagged CHA0 (CHA0-*gfp2*) strain was provided by Prof. Christoph Keel (The Department of Fundamental Microbiology at the University of Lausanne). The GFP-tagged Pto DC3000 (DC3000-*gfp2*) strain was provided by Prof. Zhaojun Ding (The Key Laboratory of Plant Cell Engineering and Germplasm Innovation, Ministry of Education, College of Life Sciences, Shandong University).

### Rhizosphere microbiome transplant assay

The rhizosphere microbiome transplant experiment was performed as described previously with minor modifications ^33^. First, Col-0, *fer-4*, *fer-4/FER^ΔJM^-GFP*, *fer-4/FERN-GFP*, *fer-4/FERN-GFP-NES*, *at2-mmp-1* and *At2-MMP-*OE #2 were grown for 4 weeks (5 plants per pot) in natural soil to allow assembly of a genotype-specific rhizosphere microbiome. Then, the shoots were cut and all soil from the same genotype was thoroughly mixed together in a sterilized container. That same day, the mixed soil (with genotype-specific microbiomes) was placed into new clean pots and planted with Col-0 plants. The trays and growth chamber were sterilized with 70% ethanol before the experiment and all plants were watered with autoclaved water. Different genotypes were put in separate trays side by side in the same growth chamber and grown under 80-100 μE light on a 16-h light/8-h dark cycles. Only non-bolting plants were chosen for the rhizosphere sampling to avoid the effects of differences in developmental stage.

### Bacterial root inoculation assays

To inoculate roots with DC3000-*gfp2* or CHA0-*gfp2*, spore suspensions were dropped onto solid half-strength MS medium without sucrose following a previously described method ^9^ with some modifications. In short, uniform 5-day-old seedlings were selected and transferred to half-strength MS medium solidified with 1% agar without sucrose (pH 5.8). After incubation overnight in LB medium, bacteria were collected, washed and resuspended in distilled water. Then, 10 μL of bacterial suspension at an optical density of OD_600_ = 0.1 for DC3000-*gfp2* or OD_600_ = 0.02 for CHA0-*gfp2* was applied by depositing small droplets along the whole root. Infected seedlings were then grown vertically for 3 days before microscope observation. The bacterial solution was carefully removed from the roots by washing with PI solution for 1 min and roots were transferred to sterile MiliQ water for 5 s before confocal imaging.

### Colony forming units (CFU) counting

The CFU counting method was used to quantify the level of CHA0-*gfp2* colonization^9^. Briefly, 5-day-old seedlings were inoculated by the drop-inoculation method described above. 3 days post-inoculation, Col-0, *fer-4*, *fer-4/FER^ΔJM^-GFP*, *fer-4/FERN-GFP*, *fer-4/FERN-GFP-NES*, *at2-mmp-1*, and *At2-MMP-*OE #2 seedlings were imaged, then harvested, gently washed by dipping them in sterile deionized water, dried on sterile filter paper and collected in sterile Eppendorf tubes containing 500 μL of extraction buffer (10 mM MgCl_2_ and 0.01% [v/v] Silwet L-77). Three seedlings were pooled together for each of the three technical replicates. Seedlings were homogenized using TissueLyser II (QIAGEN, Germany) with stainless steel beads. Samples were diluted in series in 10-fold steps, then 20 µL of each solution was spotted onto LB agar plates. CFUs were counted for the most appropriate dilution (final dilution: 4,000- or 10,000-fold) after 24 h incubation at 28℃ until colonies were clearly visible. The total CFU was normalized by centimeters of root length measured from images ([CFU/mL × 500/3]/mean root length). The experiment was conducted in three biological replicates.

### Quantification and statistical analysis

For quantifying the nucleus-localized fluorescence intensity of MAMP markers, confocal images were analyzed with the Fiji package (http://fiji.sc/Fiji). Contrast and brightness were adjusted in the same manner for all images. All statistical analyses were done with Graphpad Prism 8.0 software. One-way analysis of variance (ANOVA) was performed, and Tukey’s test was subsequently used as a multiple comparison procedure. Details about the statistical approaches used can be found in the figure legends. The data are presented as means ± standard deviation (SD), and ‘‘n’’ represents the number of plant roots.

---

**Extended Data Fig. 1.**
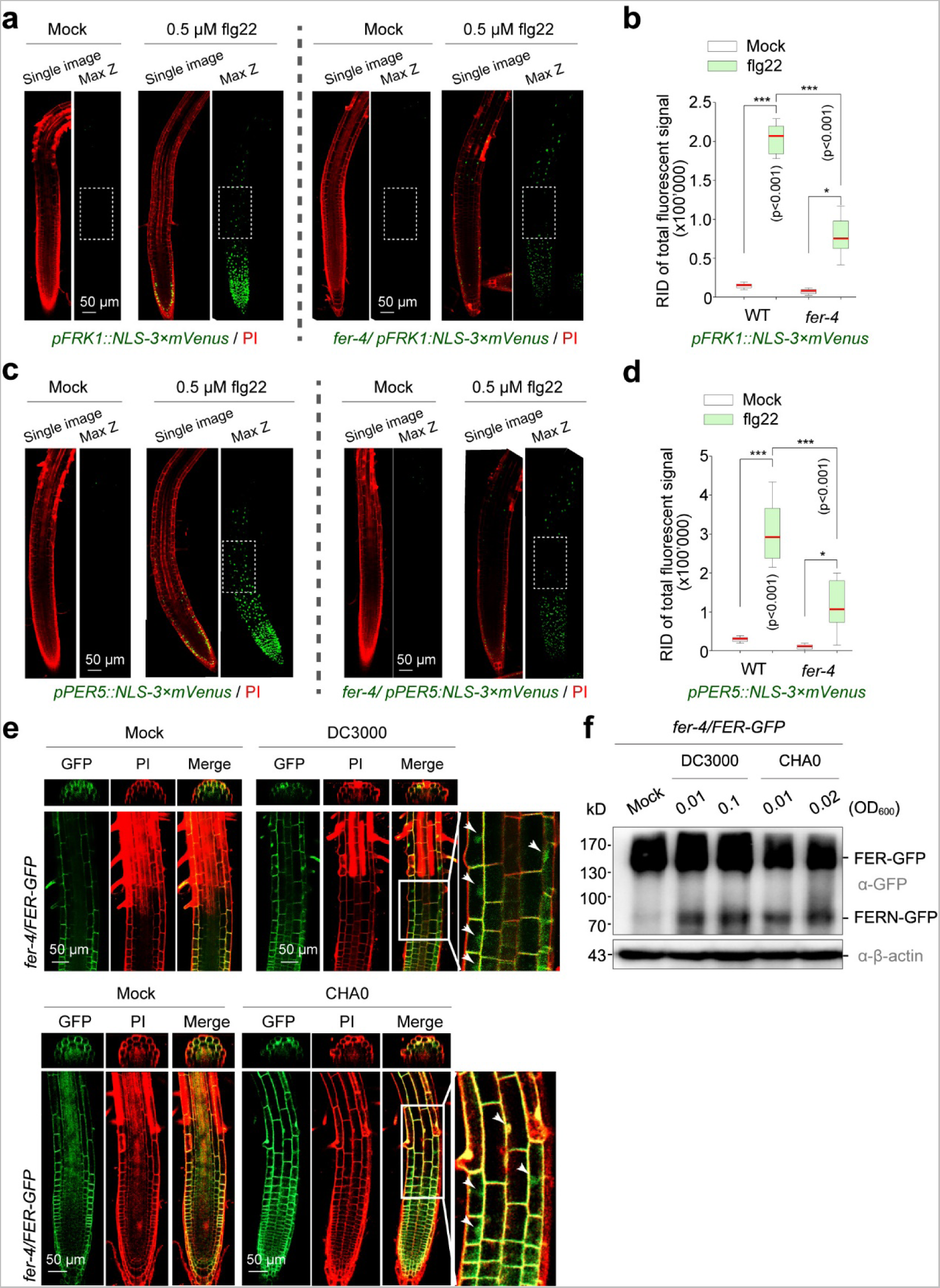
FER mediates immune responses in the TZ and EZ of roots. (**a**) Expression patterns of MAMP promoter marker line (*pFRK1*) in response to 0.5 μM flg22 for 8 h in Col-0 and *fer-4* background. mVenus signals (green) co-visualized with propidium iodide (PI, red). The white dashed box marks the TZ and EZ of roots. (**b**) Quantitative analysis of mVenus signal intensities of the *FRK1* marker in the absence or presence 0.5 μM flg22. RID, raw intensity density. Boxplot centers show median (n = 12 roots). Asterisks (*p < 0.05, ***p < 0.001) indicate statistically significant differences between means by ANOVA and Tukey’s test analysis. (**c**) Expression patterns of MAMP promoter marker line (*pPER5*) in response to 0.5 μM flg22 for 8 h in Col-0 and *fer-4* background. The white dashed box marks the TZ and EZ of roots. (**d**) Quantitative analysis of mVenus signal intensities of the *PER5* marker in the absence or presence 0.5 μM flg22. Boxplot centers show median (n = 12 roots). Asterisks (*p < 0.05, ***p < 0.001) indicate statistically significant differences between means by ANOVA and Tukey’s test analysis. (**e**) GFP fluorescence localization in D-roots of *fer-4/FER-GFP* in response to DC3000 at OD_600_ = 0.1 or CHA0 at OD_600_ = 0.02 treatments for 8 h in D-roots. GFP signals (green) co-visualized with propidium iodide (PI, red). Single confocal microscope images are presented. (**f**) Immunoblotting of FER cleavage in *fer-4/FER-GFP* seedlings in response to DC3000 or CHA0 for 8 h in D-roots.

**Extended Data Fig. 2.**
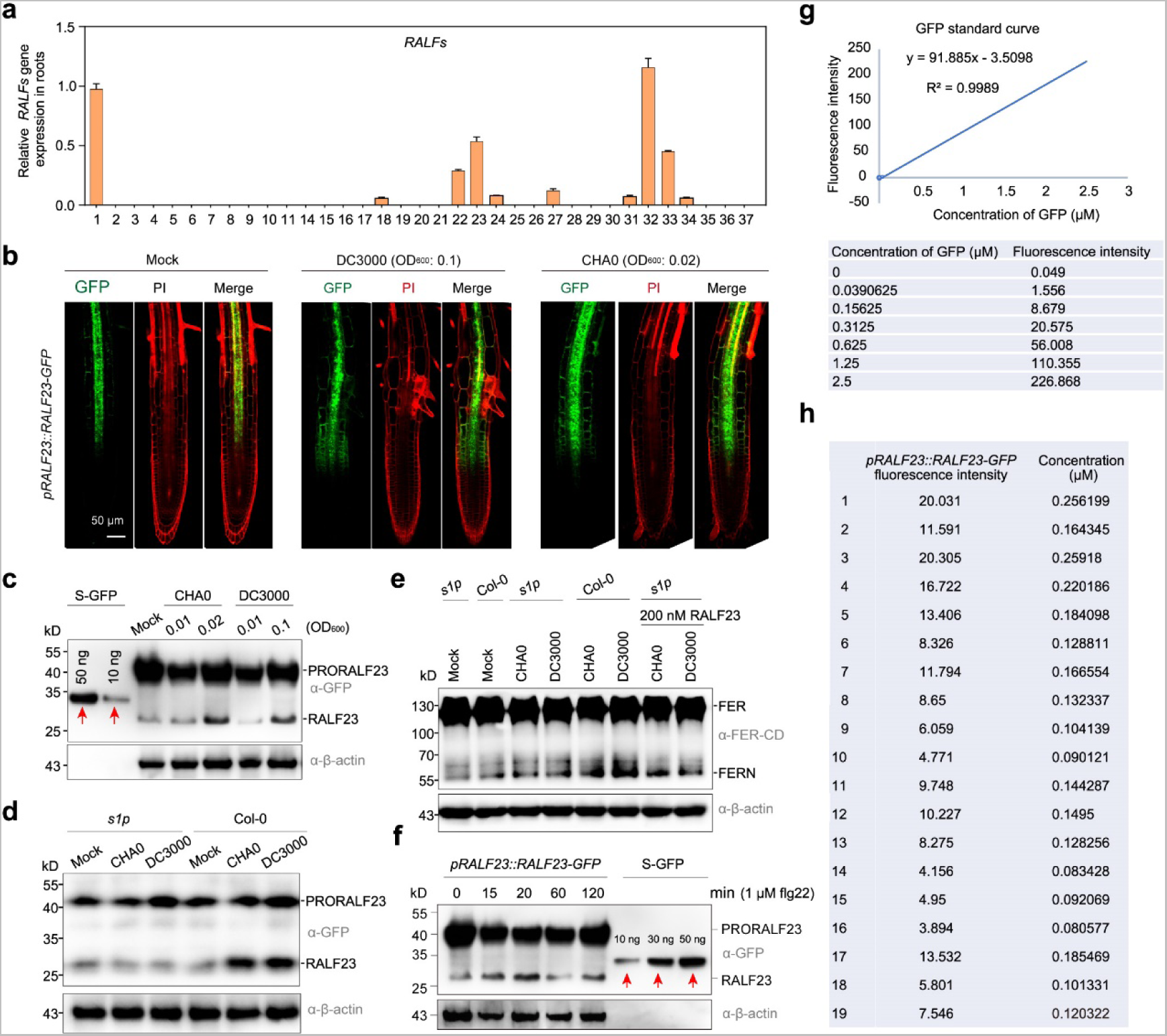
The role of RALF23 in the immune response in roots and the physiological concentration analysis of mature RALF23. (**a**) Root-expressed *RALF* genes. Col-0 seedlings were grown on solid half-strength MS plates for 5 days before their roots were collected for RNA extraction. Data are means ± SD, *n* = 3. *ACTIN2* was used as an internal control, and all values are relative to *RALF1* (set as 1.0). (**b**) Fluorescence patterns in the *pRALF23::RALF23-GFP* seedlings in response to DC3000 or CHA0 for 8 h in D-roots. Images show maximum projection of Z-stacks imaging MZ, TZ and EZ. Photographs were taken with similar settings. (**c**) Estimation of the concentration of mature RALF23 in *pRALF23::RALF23-GFP* seedlings. The red arrows represent GFP standard samples. (**d**) Cleavage of RALF23 in Col-0 or *s1p-6* mutant seedling in response to DC3000 or CHA0 for 8 h in D-roots. (**e**) Cleavage of FER in Col-0 or *s1p* mutant seedling in response to DC3000 or CHA0, or in *s1p* mutant seedling in response to DC3000 or CHA0 with 200 nM RALF23 for 8 h in D-roots. Antibody α-FER-CD was used immunoblotting. (**f**) Estimation of the concentration of mature RALF23 in *pRALF23::RALF23-GFP* seedlings. The red arrows represent the GFP standard samples. (**g**) Standard curve of GFP fluorescence intensity vs. concentration. The maximum molar concentration of the GFP standard curve was approximately 2.5 μM. (**h**) Fluorescence intensity of GFP in single cell of *pRALF23::RALF23-GFP* seedling roots. The concentration of RALF23-GFP was estimated using the GFP standard curve.

**Extended Data Fig. 3.**
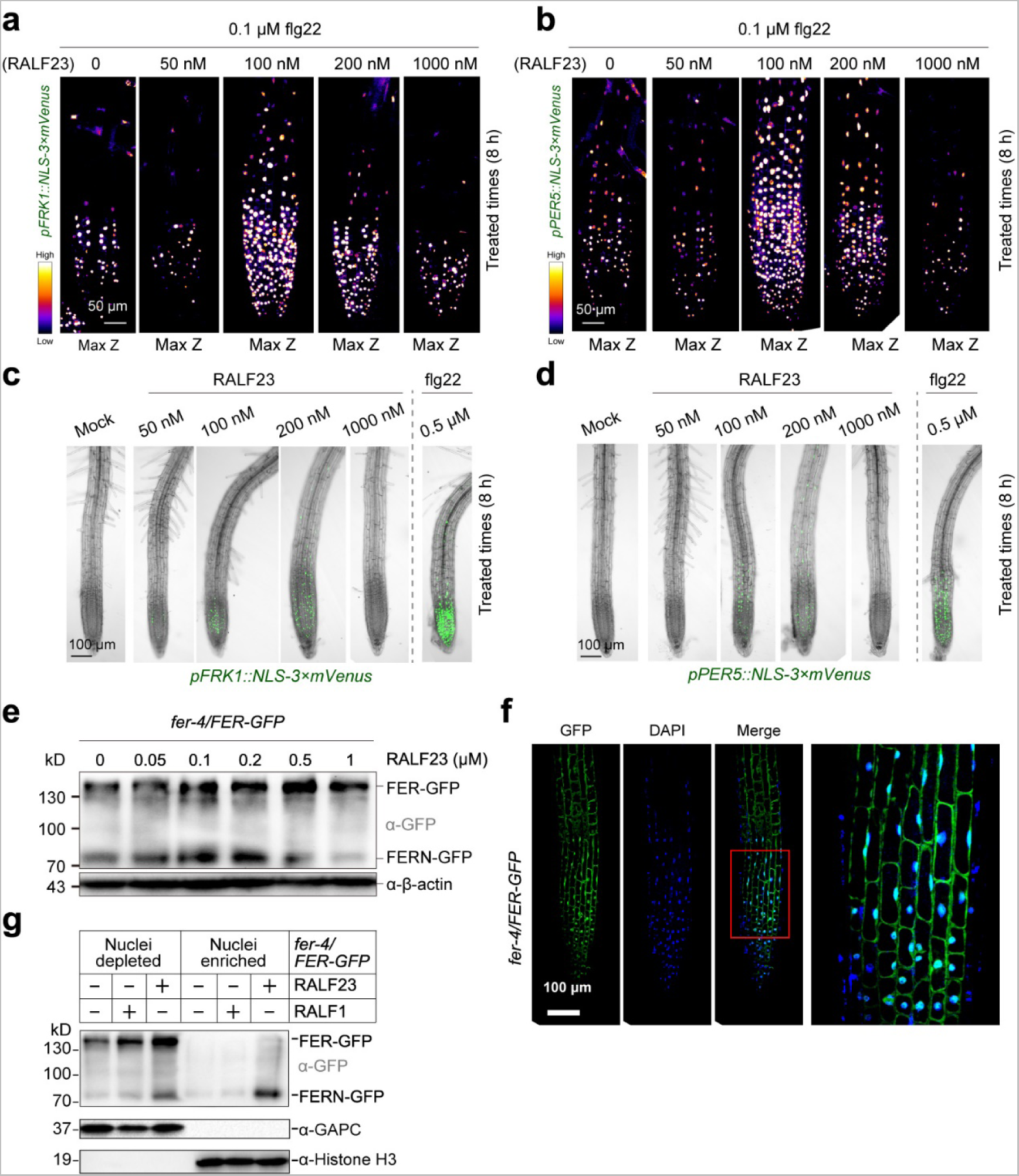
The role of RALF23 in the immune response and FER cleavage and nuclear localization in roots. (**a–b**) Fluorescence patterns in the *pFRK1::NLS-3×mVenus* (**a**) and *pPER5::NLS-3×mVenus* (**b**) MAMP promoter marker lines in response to different concentrations of RALF23 with 0.1 μM flg22 for 8 h. Images show maximum projection of Z-stacks imaging MZ, TZ and EZ. Photographs were taken with similar settings. Scale bar, 50 μm. (**c–d**) Fluorescence patterns in *pFRK1::NLS-3×mVenus* (**c**) and *pPER5::NLS-3×mVenus* (**d**) MAMP promoter marker lines in response to 50 nM to 1 μM RALF23 or 0.5 μM flg22 for 8 h. (**e**) *fer-4/FER-GFP* seedlings treated with different concentrations of RALF23. The C-terminal fragments of the FER-GFP fusion were detected using antibody α-GFP. (**f**) *fer-4/FER-GFP* roots treated with 200 nM RALF23 for 3 h stained with DAPI. All experiments were performed more than 3 times with similar results. (**g**) Immunoblot analysis of FER cleavage. Nuclei-depleted and nuclei-enriched fractions from 5-day-old *fer-4/FER-GFP* seedlings treated with RALF1 or RALF23 for 3 h were used. GAPDH and nuclear histone H3 were used as cytosolic and nuclear markers, respectively.

**Extended Data Fig. 4.**
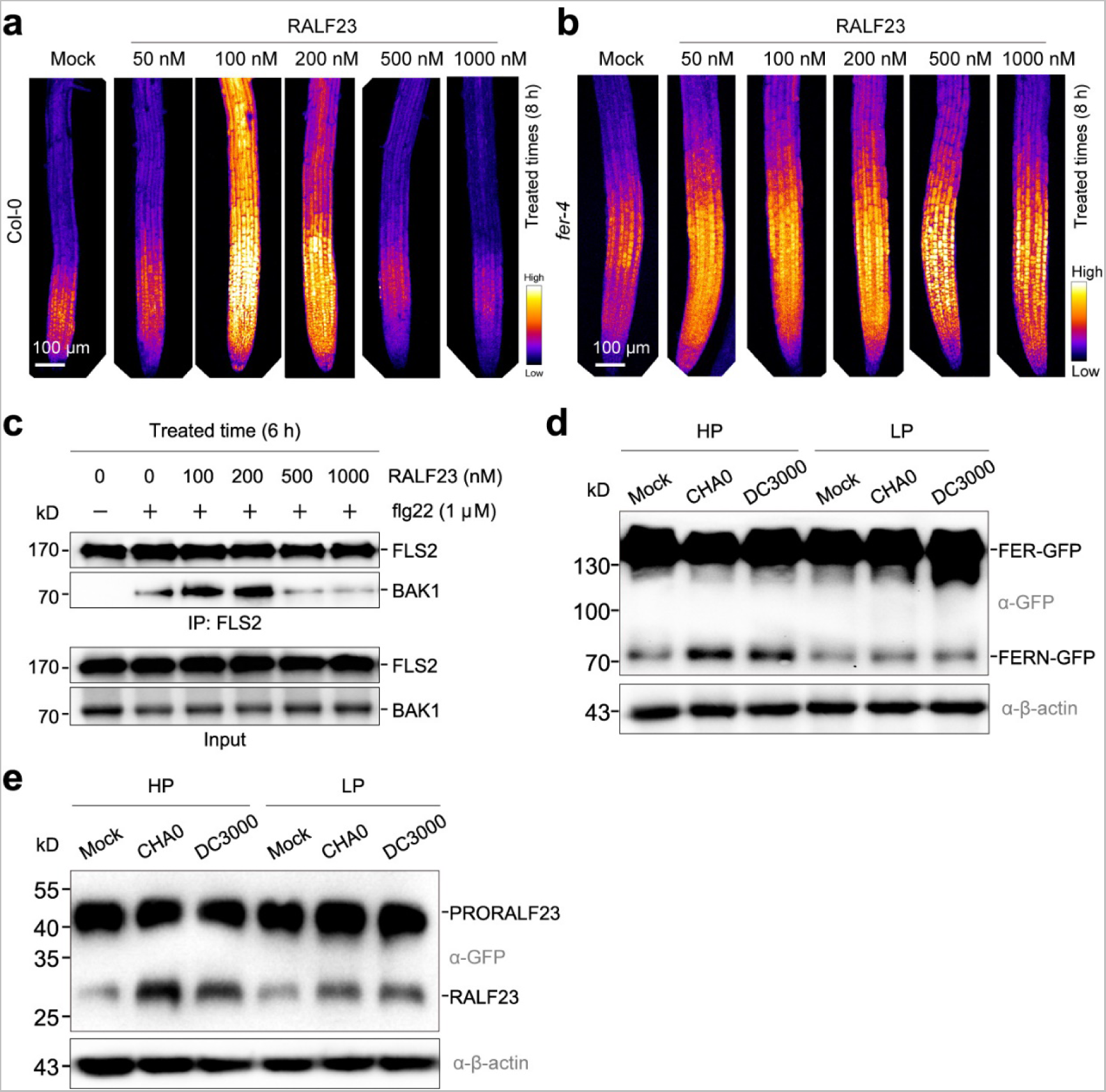
RALF23 at different concentrations has opposing effects on root immunity. (**a–b**) Representative images of H_2_DCFDA-stained roots of Col-0 (**a**) and *fer-4* (**b**) pre-treated with different concentrations of RALF23 for 3 h. Representative images from one experiment are shown, repeated three with consistent results. (**c**) RALF23 at different concentrations has opposing effects on FLS2-BAK1 complex formation. Col-0 seedlings were grown in half-strength MS media for 5 days, then treated with 1 μM flg22 alone or 1 μM flg22 in combination with varying concentrations of RALF23 (100, 200, 500, 1000 nM) for 6 h. The amount of immunoprecipitated FLS2 and coimmunoprecipitated BAK1 were determined using anti-FLS2 and anti-BAK1 antibodies, respectively. (**d**) *fer-4/FER-GFP* seedlings grown in either HP or LP media were treated with DC3000 or CHA0 for 8 h in D-roots. (**e**) *pRALF23::RALF23-GFP* seedlings grown in either HP or LP media were treated with DC3000 or CHA0 for 8 h in D-roots. Antibody α-GFP was used immunoblotting.

**Extended Data Fig. 5.**
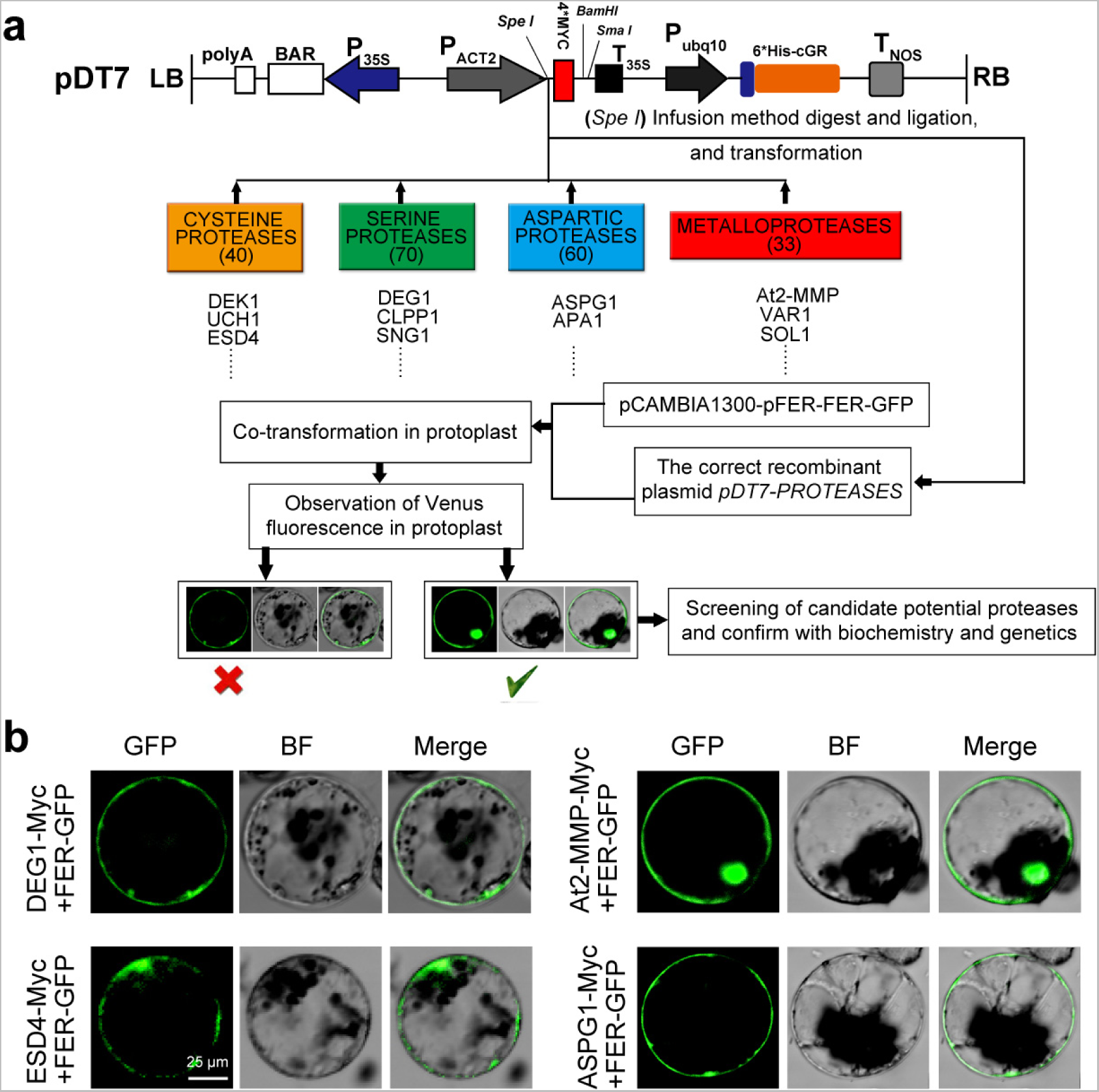
Screening of proteases cleaving FER. (**a**) Schematic diagram of protease gene overexpression based on pDT7. A total of 203 protease genes predicted by bioinformatics to be expressed in Arabidopsis roots, belonging to four gene subfamilies, were individually introduced into pDT7 and then transformed together with the *pFER::FER-GFP* construct into root protoplasts. The proteases that caused nucleus-localized GFP fluorescence in GFP-positive protoplasts were considered candidates that may induce FER cleavage. (**b**) Representative images of protoplasts co-transformed with protease genes and *FER-GFP*.

**Extended Data Fig. 6.**
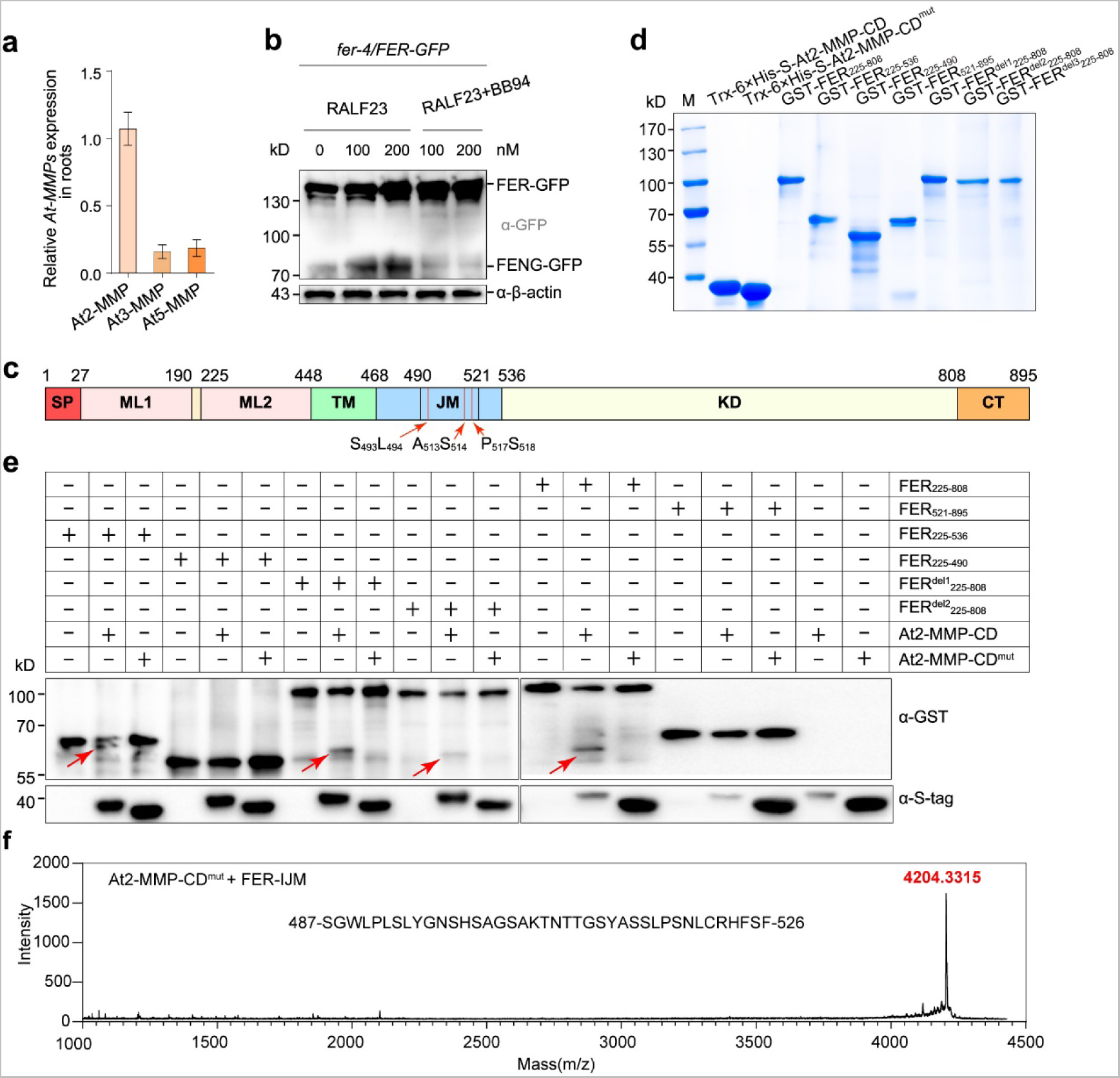
At2-MMP cleaves FER. (**a**) RT-qPCR analysis of *At2-MMP*, *At3-MMP* and *At5-MMP* expression in wild-type D-roots. *ACTIN* was used as the reference gene. Values are means ± SD, *n* = *4*. (**b**) At-MMPs cleave FER. Seedlings were treated with RALF23, or RALF23 + 5 μM BB94 for 2 h. Proteins were detected using α-GFP antibody. (**c**) A domain map based on FER. Numbers indicate amino acid positions. SP, signal peptide; TM, transmembrane domain; ML, malectin-like domain; JM, juxtamembrane domain; KD, kinase domain; CT, C-terminal. The red arrows represent the cleaved sites of FER. (**d**) Purification of recombinant proteins. (**e**) Immunoblot analysis of the proteolytic activity of recombinant At2-MMP and At2-MMP^mut^ toward GST-FER_225–536_, GST-FER_225–808_, GST-FER_521–895_, FER^del1^_225–808_ and FER^del2^_225–808_. S_493_L_494_, A_513_S_514_ and P_517_S_518_, named FER^del3^_225–808_; S_493_L_494_, named FER^del1^_225–808_; and S_493_L_494_ and P_517_S_518_, named FER^del2^_225–808_. The red arrows represent the cleaved N-terminal of GST-FER. (**f**) MALDI-TOF MS analysis of FER-IJM cleaved by At2-MMP-CD^mut^.

**Extended Data Fig. 7.**
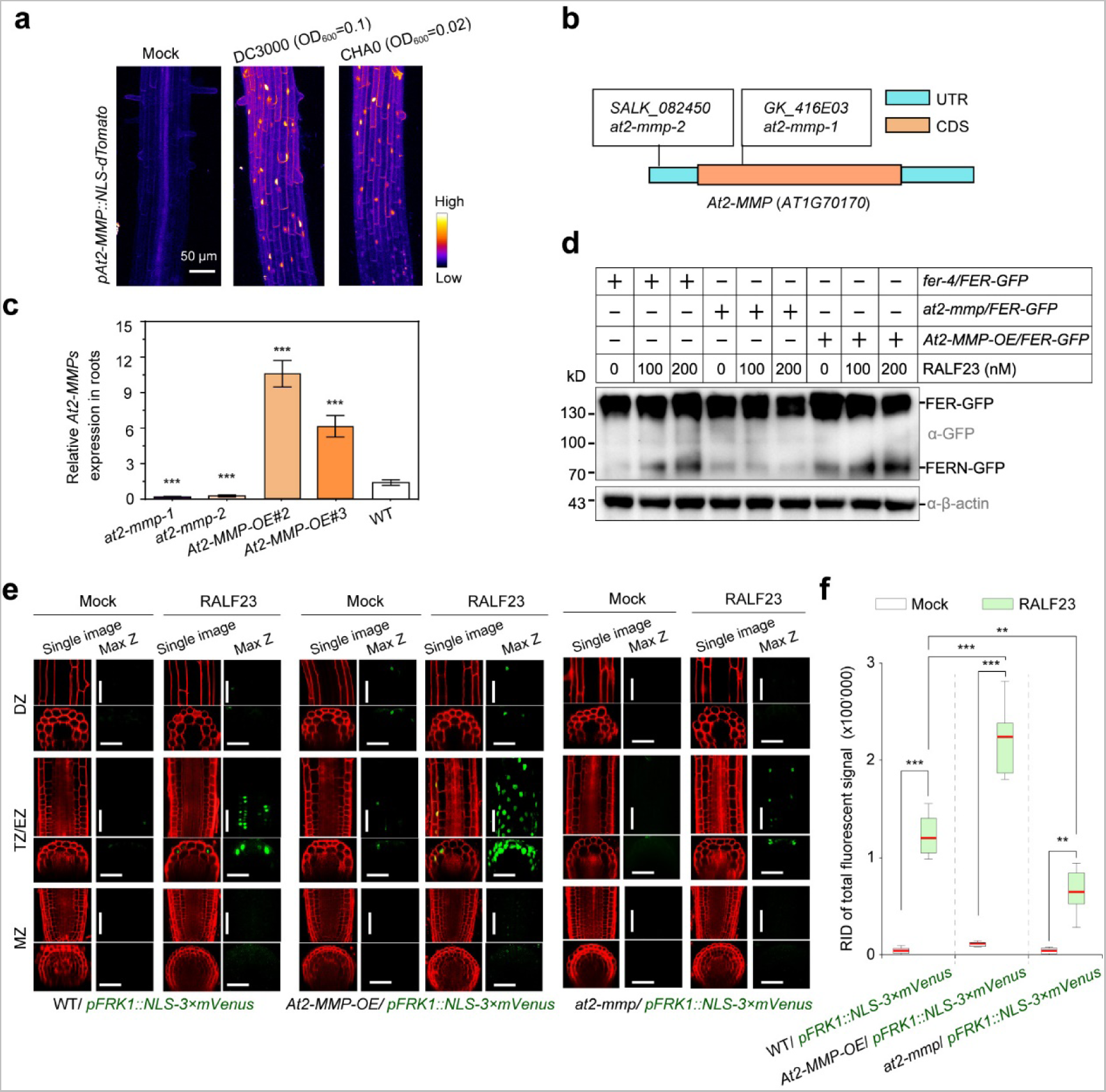
At2-MMP cleaves FER, mediating RALF23-driven localized immune response. (**a**) Fluorescence patterns in the *pAt2-MMP::NLS-dTomato* seedlings in response to DC3000 or CHA0 for 8 h in D-roots. Images show maximum projection of Z-stacks imaging TZ and EZ. Photographs were taken with similar settings. (**b**) Identification of *at2-mmp* mutants. Schematic diagram showing the T-DNA insertion sites of the *at2-mmp-1* (GK_416E03) and *at2-mmp-2* (SALK_082450) single knockout lines in the Col-0 background. (**c**) Relative expression of *At2-MMP* in the roots of different transgenic lines. *At2-MMP* expression levels were quantified relative to *ACTIN* levels. Values are means ± SD, *n* = *3*. Student’s *t* test. ***p < 0.001. (**d**) At2-MMP cleaves FER in plants. *fer-4/FER-GFP*, *at2-mmp-1* and *At2-MMP-*OE#2 in the *fer-4/FER-GFP* background were treated with RALF23 for 3 h or with H_2_O as mock control. (**e**) mVenus fluorescence pattern of the MAMP promoter marker line *pFRK1::NLS-3×mVenus* in response to 200 nM RALF23 for 8 h in the Col-0, *At2-MMP-*OE#2, and *at2-mmp-1* backgrounds. mVenus signals co-visualized with PI. (**f**) Quantitative analysis of mVenus signal intensities of the *FRK1* marker in response to 200 nM RALF23 for 8 h at TZ/EZ in the Col-0, *At2-MMP-*OE#2, and *at2-mmp-1* backgrounds. Boxplot centers show median (n = 12 roots). Asterisks (***p < 0.001, **p < 0.01) indicate statistically significant differences between means by ANOVA and Tukey’s test analysis.

**Extended Data Fig. 8.**
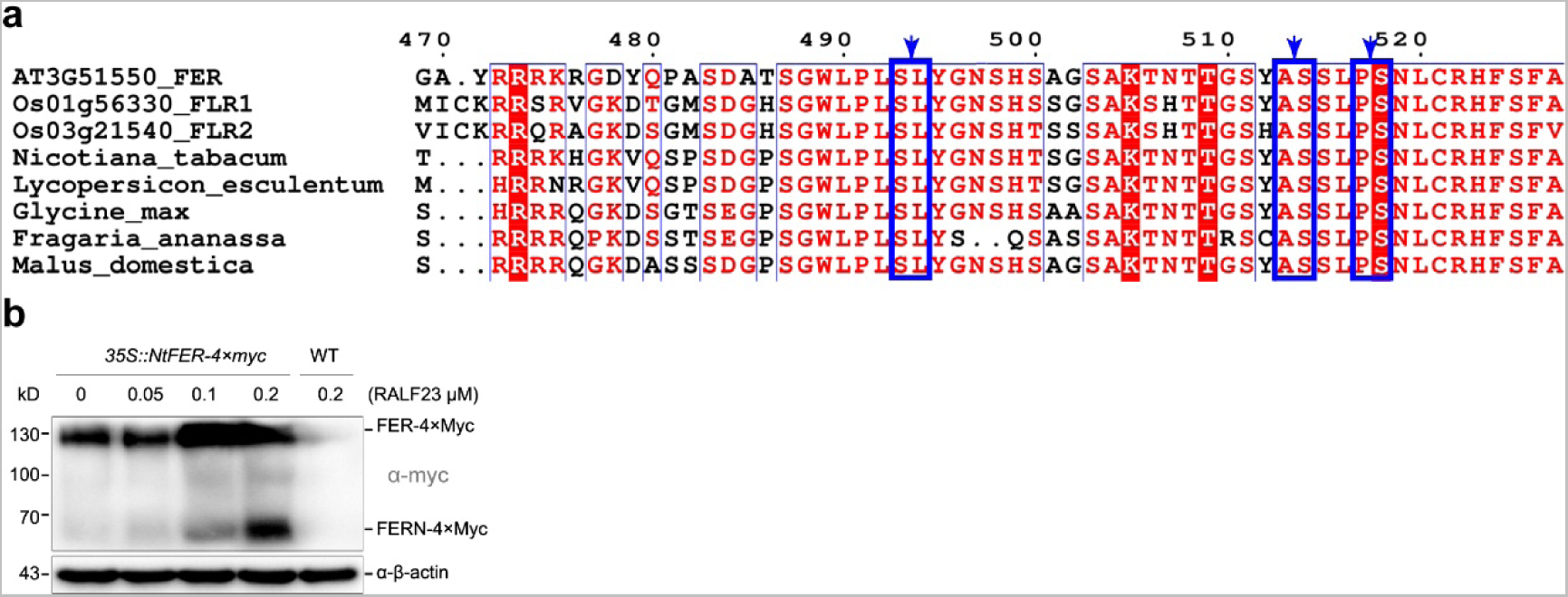
The cleavage sites of FER are conserved in multiple plant species. (**a**) Multiple sequence alignments of the intracellular juxtamembrane domains of FER in Arabidopsis and its homologs in rice (*Oryza sativa*), tobacco (*Nicotiana tabacum*), tomato (*Lycopersicon esculentum*), soybean (*Glycine max*), strawberry (*Fragaria* × *ananassa*) and apple (*Malus domestica*). The blue arrows represent conserved FER cleavage sites by At2-MMP. (**b**) Tobacco FER (*Nt*FER) cleavage in *35S::NtFER-4×Myc* seedlings in response to RALF23 for 3 hours. The C-terminal protein fragments were detected using α-myc antibody.

**Extended Data Fig. 9.**
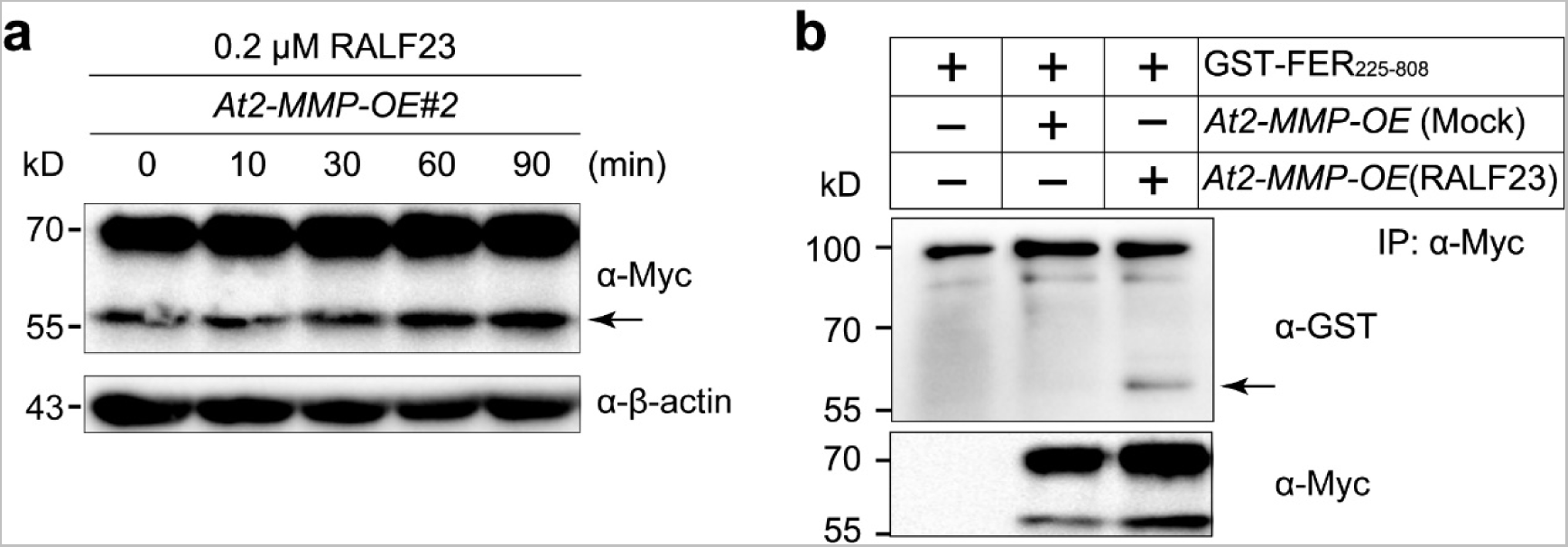
RALF23 regulates At2-MMP activity. (**a**) RALF23 induces the accumulation of low-MW At2-MMP. The arrow represents the low-MW At2-MMP. (**b**) Immunoblot analysis of the proteolytic activity of At2-MMP induced by RALF23 toward FER_225–808_. *At2-MMP-*OE seedlings were treated with 200 nM RALF23 or H_2_O for 3 h, then immunoprecipitated with α-Myc beads, then the immunoprecipitated productions were incubated with FER_225–808_ in cleavage buffer. The arrow represents FER cleaved by low-MW mature MMP.

**Extended Data Fig. 10.**
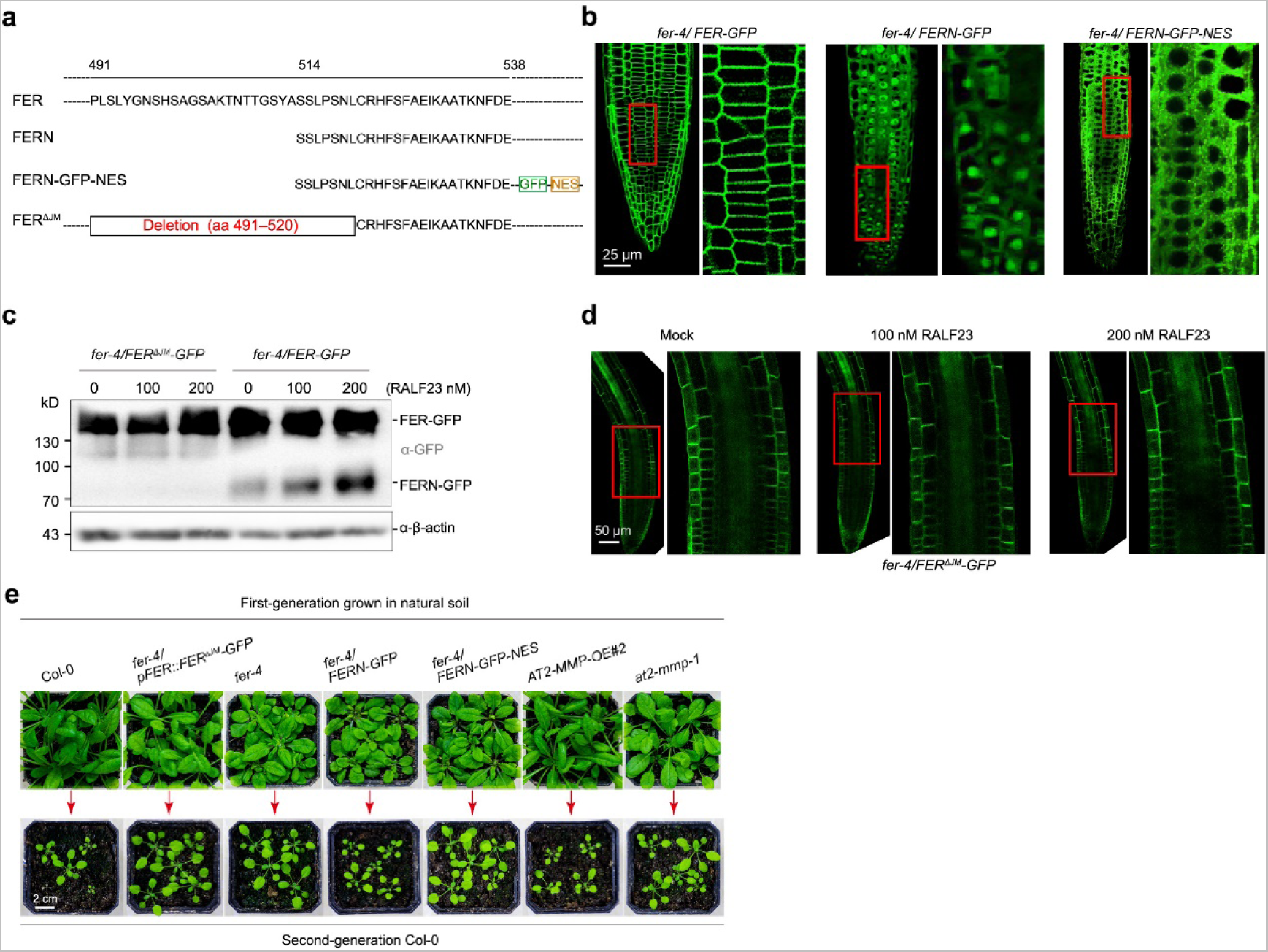
Identification and phenotypic analysis of transgenic lines of *FERN-GFP*, *FERN-GFP-NES* and *FER^ΔJM^-GFP*. (**a**) Schematic diagram of the full-length, FER^ΔJM^ and C-terminal (FERN) domain of FER. (**b**) Subcellular localization of FER and FERN in *fer-4/FER-GFP*, *fer-4/FERN-GFP* and *fer-4/FERN-GFP-NES* lines in 5-day-old D-roots. (**c**) *fer-4/FER^ΔJM^-GFP* and *fer-4/FER-GFP* seedlings in response to 0, 100 or 200 nM RALF23 for 3 hours in D-roots. The C-terminal protein fragments were detected using an α-GFP antibody. (**d**) GFP fluorescence localization of *fer-4/FER^ΔJM^-GFP* in roots after treated with RALF23 for 3 h. At least four biological replicates were performed for each assay with similar results. (**e**) Representative images of different plants grown in natural soil (generation 1) and wild-type Col-0 plants (generation 2).

**Table S1.**
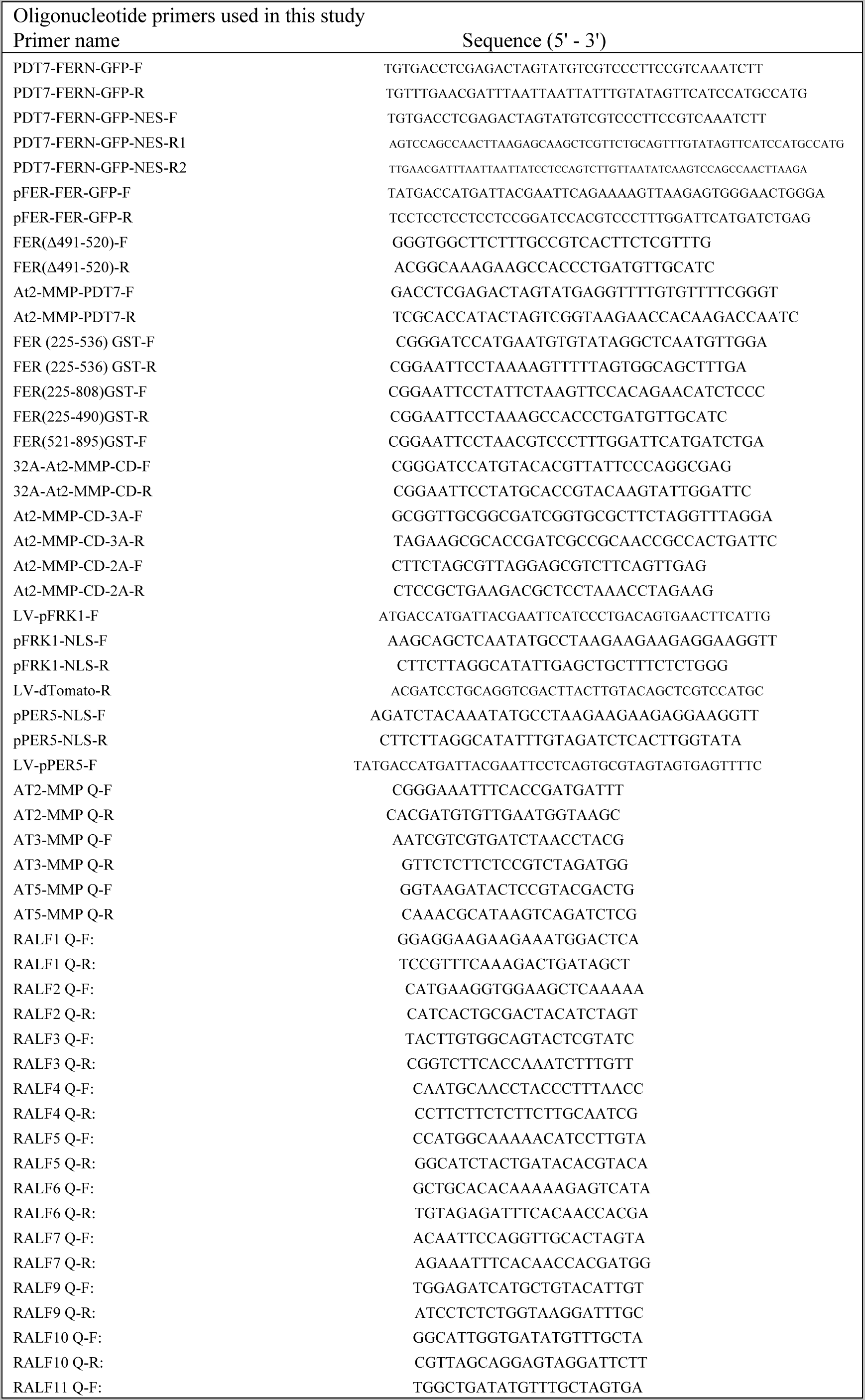
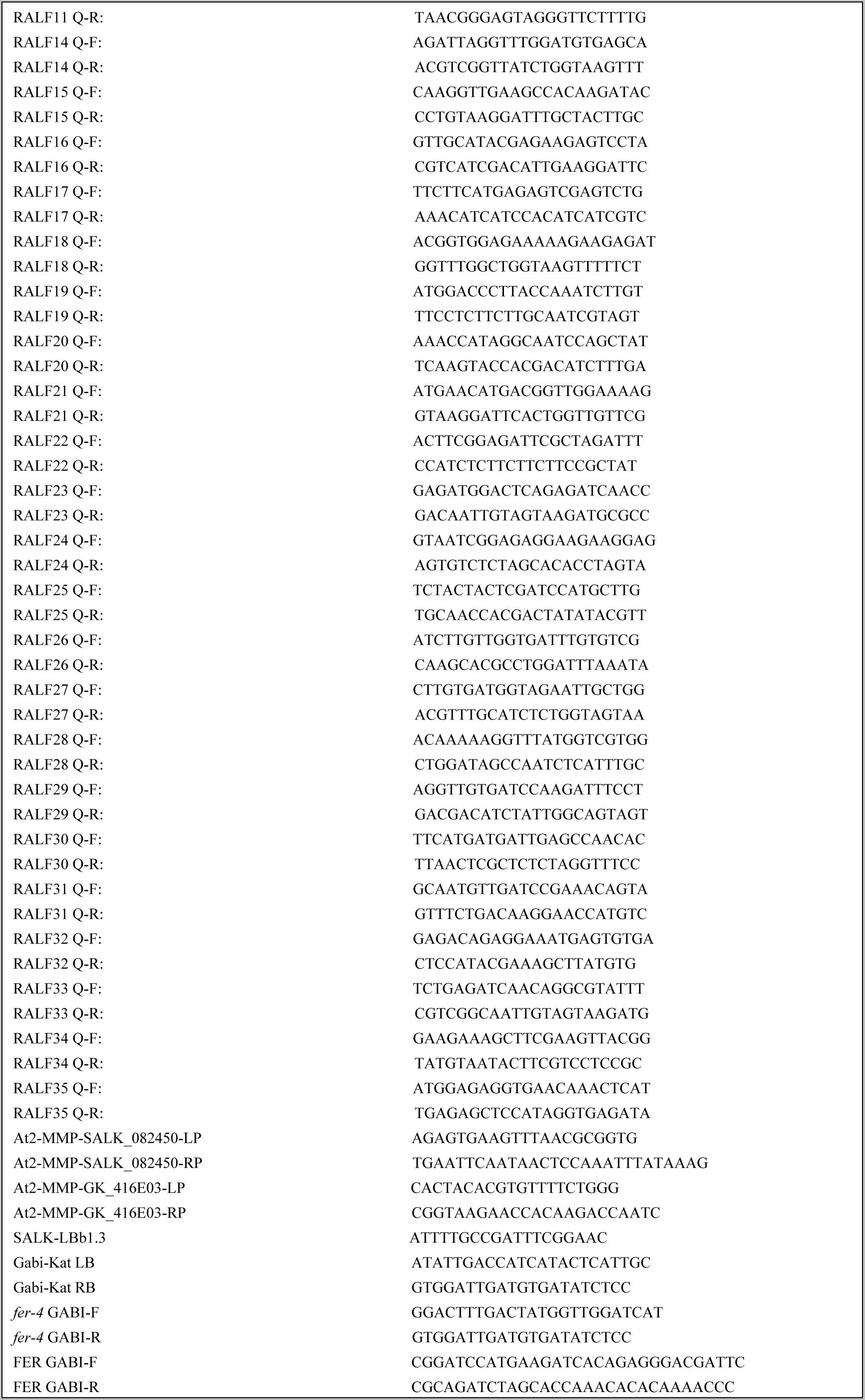

